# Unravelling the ancient fungal DNA from the Iceman’s gut

**DOI:** 10.1101/2024.01.24.576930

**Authors:** Nikolay Oskolkov, Anna Sandionigi, Anders Göterström, Fabiana Canini, Benedetta Turchetti, Laura Zucconi, Tanja Mimmo, Pietro Buzzini, Luigimaria Borruso

**Affiliations:** Department of Biology, National Bioinformatics Infrastructure Sweden, Science for Life Laboratory, Lund University, Lund, Sw Anna Sandionigi eden; Department of Informatics, Systems and Communication, University of Milan-Bicocca, Milan, Italy; Quantia Consulting srl, Milan, Italy; Centre for Palaeogenetics, Department of Archaeology and Classical Studies, Stockholm University; Department of Ecological and Biological Sciences, University of Tuscia, 01100 Viterbo, Italy; Department of Agricultural, Food and Environmental Sciences, University of Perugia, Perugia, Italy; Faculty of Agricultural, Environmental and Food Sciences, Free University of Bozen-Bolzano, Piazza Università 5, Italy

**Keywords:** fungi, ancient DNA (aDNA), Pseudogymnoascus Iceman

## Abstract

Here, we explore the possible ancient fungal species in the gut of Ötzi, the Iceman, a naturally mummified human found in the Tyrolean Alps (border between Italy and Austria). While ancient DNA (aDNA) has been extensively used to study human, animal, and plant evolution, this research focuses on ancient microbial diversity, specifically fungi. Fungal DNA is often underestimated in metagenomic samples, however here we hypothesise the possibility of retrieving ancient fungal sequences from Ötzi’s gut. A robust bioinformatic pipeline has been developed to detect and authenticate fungal aDNA from stomach, small intestine, and large intestine samples. We revealed the presence of *Pseudogymnoascus* genus, with *P. destructans* and *P. verrucosus* as possible species, that were particularly abundant in the stomach and small intestine. We suggest that Ötzi may have consumed these fungi accidentally, likely in association with other elements of his diet, and they thrived in his gut after his death due to their adaptability to harsh and cold environments. This research provides insight into the coexistence of ancient humans with specific fungal species and proposes and validates a conservative bioinformatic approach for detecting fungal aDNA in historical metagenomic samples.

**Significance statement:** Despite their essential interactions with all kingdoms of life, limited molecular studies have focused on ancient fungi. Here, we developed a thorough bioinformatic pipeline that allowed us to detect the presence of ancient DNA likely belonging to *Pseudogymnoascus destructans* and *P. verrucosus* in the gut of Ötzi, a human naturally mummified over 3,000 years ago in the Tyrolean Alps. Both species can survive harsh environmental conditions, and *P. destructans* is known for its pathogenicity, suggesting that Ötzi may have accidentally ingested them and providing valuable insights into how ancient humans coexisted with specific fungal species. We propose a highly reliable methodology for detecting ancient fungal DNA in metagenomic studies of historical samples that can have broader applications to understand ancient ecosystems and their interactions.

## Introduction

Palaeobiological approaches based on ancient DNA (aDNA) have extensively been used to study the evolution of humans, animals, and plants (1–5). Recent advances in sedimentary and environmental ancient metagenomics provided unprecedented time series resolution for understanding the evolution of flora and fauna without macroscopic remains such as bones or teeth (6–9). Studies based on whole shotgun metagenomic sequencing leverage exciting information on bacterial species present in ancient samples (10–12). Nevertheless, several challenges complicate ancient microbial metagenomics analysis, such as modern contamination, which makes authentication analysis of major importance (13, 14). In addition, several different microbial taxa are typically present in metagenomic samples (15). Thus, studying ancient microbiomes, especially when it comes to fungi, remains a challenge. Fungal DNA is often underrepresented in metagenomic samples due to several limitations of sequencing methods (16). According to the last estimates, the kingdom of fungi includes up to twelve million species (17), that play pivotal ecological roles in all ecosystems (e.g. in soil or dead material) as saprotrophs and in association (e.g., as symbionts or parasites) with plants, animals and humans, including their gut microbiome (18, 19). In the last few years, several studies have demonstrated that fungi are crucial components of the gut microbiome (20). Nevertheless, the gut microbiome is dramatically susceptible to the environment; thus, distinguishing between non-resident fungal species ingested due to diet or, more generally, acquired by the environment and resident taxa is challenging (19, 21).

Here, we hypothesised the possible presence of ancient fungal species associated with the gut of the Iceman, nicknamed Ötzi, the naturally mummified human body found in 1991 in the Similaun glacier at 3,200 m asl in the Tyrolean Alps (border between Italy and Austria) (22) (Figure 1).

**Figure 1.**
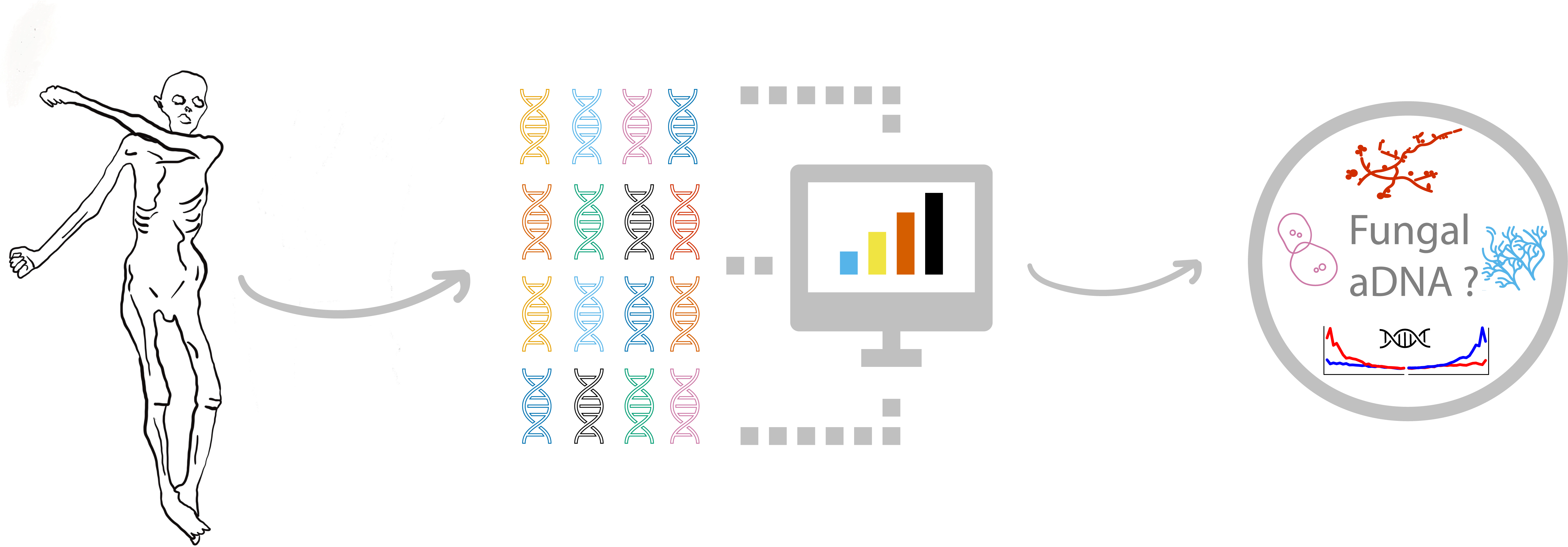
Graphical abstract of workflow about the ancient fungal DNA detection in iceman sample.

Some works have already demonstrated the presence of aDNA fragments of a few bacterial species in the Iceman’s gut (12, 23), but to the best of our knowledge, there is no information regarding the possible presence of ancient fungi. To test our hypothesis, we applied a robust bioinformatic pipeline to detect the endogenous genetic signals of fungal aDNA by using sequence data from the stomach, small intestine, and large intestine and considering muscle tissues as the negative control (Figure 2). In addition, a blank sample was used as a second negative control.

**Figure 2.**
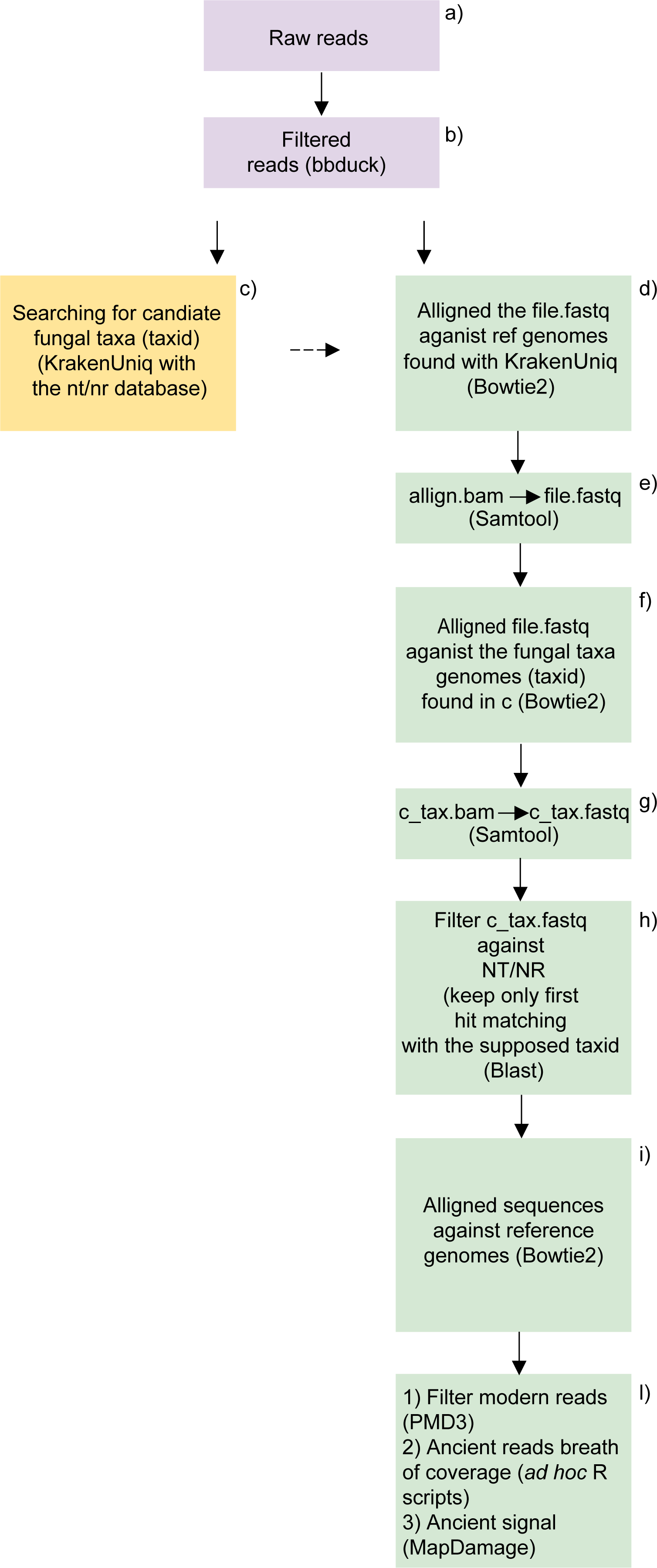
Flowchart of the bioinformatic pipeline used to analyse the Iceman fungal community.

## Methods

The Iceman dataset analysed in this study was downloaded from the European Nucleotide Archive (ENA) under accession number ERP012908, all information about the samples and sequencing details can be found in Maixner et al (12). Raw sequences were analyzed using a bioinformatic pipeline consisting of three main phases: i) quality control, ii) taxonomic identification of the fungal taxa, and iii) authentication analysis of the detected fungi. All pipeline steps are illustrated in the flowchart of Figure 2. The software BBDuk and BBMerge (24) were used to trim Illumina adapters and merge overlapping pair-end metagenomic reads. FastQC was used for inspecting the quality metrics of the pre-processed reads (25). Next, trimmed, and merged reads were taxonomically classified via a k-mer based approach with KrakenUniq (26) to get an overview of the microbial community in the samples and retrieve a fast taxonomic assignment for each read. Sequencing data originating from the organisms identified by KrakenUniq were conservatively filtered by depth (min 200 reads) and breadth of coverage (min 1000 unique k-mers), which aimed to reduce the number of false-positive discoveries. Further, for unbiased and robust taxonomic classification with KrakenUniq, we used one of the largest and most diverse nucleotide databases, which is NCBI GenBank (NT), accessed in December 2020, that included all microbial, vertebrate, non-vertebrate, and plant organisms, specifically 73,300,584 reference sequences corresponding to 9,556,710 taxa with 1,855,732 species. The KrakenUniq NCBI GenBank nucleotide (NT) database is publicly available through the SciLifeLab Figshare at https://doi.org/10.17044/scilifelab.20205504. An organism was considered “detected” by KrakenUniq if it had at least 200 reads and at least 1000 unique k-mers assigned to it. The former quality metric implies filtering for depth of coverage, and the latter quality metric corresponds to the breadth of coverage, i.e. the reads have to uniformly cover a fungal reference genome for the fungus to be considered to be present in a sample. Further, we excluded the microorganisms found in the blank negative control samples from the final list of microbes detected by KrakenUniq. Hence, we did not proceed with validation and authentication analyses of these microbes.

Complementary to the k-mer based classification with KrakenUniq, the pre-processed reads were also aligned to NCBI GenBank (NT) with Bowtie2 (29) (the database is publicly available via SciLifeLab Figshare at https://doi.org/10.17044/scilifelab.21070063), which allowed for a preliminary inspection of coverage and deamination signals with MapDamage (30). Next, even if the NCBI GenBank (NT) database is optimal for organism detection because it includes a variety of organisms, it often needs to improve the quality of reference genomes to make computational analyses feasible. Therefore, for a comprehensive follow-up of an interesting candidate, it is typically required to have alignments against a good-quality reference genome. Thus, according to the taxonomic composition found with KrakenUniq, reference genomes of the most abundant fungal species were downloaded from the NCBI RefSeq resource and the pre-processed metagenomic reads were aligned again to the reference genomes by Bowtie2 (27). These new good-quality alignments were used for computing a deamination profile with MapDamage (28) and evenness of coverage with Samtools (29). Only fungal candidates that demonstrated convincing deamination and uniform evenness of coverage profiles were considered as likely present in a sample and of ancient origin. To verify KrakenUniq and Bowtie2 findings, we implemented a new alignment iteration via Blast (30). For this purpose, the reads aligned to fungal references were extracted and used for Blast analysis with the NCBI NR / NT database. Only sequences matching the first hit with the expected *taxid* were retained. To follow up the most interesting fungal candidates, further sorting of damaged reads was performed with PMDtools (31). To confirm the previously observed signals, we again evaluated the reads selected by Blast and PMDtools in terms of their deamination pattern with MapDamage (28) and evenness of coverage with Samtools (29). Finally, the organisms detected in the Iceman sample were visualized as Krona plots (32). The software Jellyfish was utilized to extract and count the k-mers (where k = 5) of both the potential ancient fungal candidates and all reference genome species belonging to their respective genera (i.e. *Pseudogymnoascus*) available in NCBI (33). Additionally, four outgroup species (*Podosphaera leucotricha* PuE-3 k121 111, *Oidiodendron maius* Zn, *Xenosphaeropsis pyriputrescens* CBS 115176, and *Drepanopeziza brunnea* ‘multigermtubi’ MB m) selected from the same taxonomic phylum (Ascomicota) and class (Leotiomycetes) of *Pseudogymnoascus*, were included, along with two species from the same phylum (*Saturnispora hagleri* NRRL Y−27828 and *Saccharomyces cerevisiae* S288C) (Ascomicota).

The matrix was normalised and log-transformed. Finally, cluster analysis based on Manhattan distance was performed with the ‘vegan’ and ‘pvclust’ R packages (34, 35).

In summary, the bioinformatic pipeline applied in this work comprised multiple complementary detection, validation and authentication steps that aimed to retrieve the most confident fungal signals. We focused only on fungal hits that resulted highly reliable concerning the choice of analysis method, i.e. we considered as highly likely true positive ancient fungal hits only those detected by both KrakenUniq, Bowtie2 and Blast and successfully authenticated by MapDamage and PMDtools. All the scripts from the bioinformatic pipeline used in this study and the main output files can be found at https://github.com/NikolayOskolkov/Iceman_fungi.

## Results and Discussion

With our analysis approach, we reproduced the main microbial findings of the Iceman reported by Maxiner et al. (11, 13), such as *Helicobacter pylori* and a few other bacteria belonging to *Clostridium* and *Pseudomonas* genera (Table SM 1; Figure SM 1 and SM 2). However, since this study focuses on the ancient fungal community of the Iceman samples, we will concentrate only on fungal hits and discuss their abundance and quality metrics (Figure SM 3). The fungal species identified by KrakenUniq and validated with conservative depth and breadth filters are presented in (Table 1). Two fungal genera were standing out as particularly abundant *Rhodotorula* with the species *R. toruloides* (19,1057 reads and 14,868 unique k-mers assigned), and *Pseudogymnoascus* with *P. destructans* (1,298,527 reads and 473,667 unique k-mers assigned), *P. verrucosus* (437,040 reads and 405,670 unique k-mers assigned) and *P. pannorum* (787,723 reads and 405,670 unique k-mers assigned) (Table 1). The assigned unique k-mers for the two genera are far above our conservative detection limit of at least 1000 unique k-mers, which implies high confidence in discovering the two fungal genera (28). Further, the read alignment procedure with Bowtie2 against the full NCBI GenBank nucleotide (NT) database confirmed that the number of reads mapped uniquely (i.e. with the mapping quality above 1) was: 2605 with *R. toruloides;* 472,871 reads with *P. destructans*; 170,837 with *P*. *pannorum* and 241,46 with *P. verrucosus* (Table 1). This indicates that mapped reads have a strong affinity towards a specific species since the reads can map to millions of other reference sequences in the NCBI Genbank nucleotide (NT) database. Additionally, the breadth of coverage yielded the following results: 472,871 reads exhibited uniform coverage across the scaffolds of the representative genome of *P. destructans* (total length > 35.8 Mb), constituting 11.33% of the entire genome (Table 1). Furthermore, 241,461 reads encompassed 9.89% of the genome of *P. verrucosus*(total length > 30.17 Mb) (Table 1). In contrast, *P. pannorum* and *R. toruloides* hits had breath coverage < 1% of the reference genomes coverage thus, we discarded both taxa from further analysis (Table 1). *P. destructans* and *P. verrucosus* had relatively high mismatches captured by their edit distance plots, which did not demonstrate strictly decreasing profiles, i.e., most reads had between 1 to 3 mismatches (S.M. Figure 4a and 4b). Deamination analysis in *P. destructans* and *P. verrucosus* showed a signal of deamination profiles with transition mutation frequencies barely reaching 4-5% at the ends of the reads (S.M. Figure 5a and 5c). The distribution of reads mapped to *P. destructans* and *P. verrucosus* demonstrated a bimodal profile with a first peak at read length of ∼45 bp and a second at ∼90 bp (Figure S.M. 5b and 5d). The combination of weak deamination, relatively high mismatch rate and bimodal read length distribution can imply that a mixture of reads was mapped to the *P. destructans* and *P. verrucosus* representative genomes. Two different reasons can explain the bimodal read length distribution and relatively weak overall read deamination: modern contamination and mis-mapping. First, DNA sequences from ancient (damaged and short) and likely modern (not damaged and long) *P. destructans* and *P. verrucosus* can be present in the Iceman samples. The second reason can be that ancient and modern reads from two or more species from the *Pseudogymnoascus* genus (not included in NCBI NT database) were mis-mapped to the *P. destructans* and *P. verrucosus* representative genomes. However, the relatively high mismatch rate observed in the edit distance plot (Figure S.M. 4a and 4b) indicates that the second reason is a more plausible explanation. Therefore, we can conclude that *Pseudogymnoascus* genus was confidently detected in the Iceman data. At the same time, it is less certain that the exact two species, i.e. *P. destructans* and *P.verrucosus*, are present in the metagenomic samples. To exclude reads potentially coming from modern contamination (first reason), we filtered reads with a PMD score above 3, which implies a high confidence damage pattern (36). Indeed, a typical authentication approach includes monitoring the fragmentation of DNA sequences and the deamination pattern of cytosine bases, which occur with a higher frequency at the extremities of the aDNA fragment (28). In addition, we considered only reads shorter than 100 bp, which were more likely to be of ancient origin than longer reads (Figure 3b and 3d) (28). This filtering step significantly improved the clarity of the deamination signal, where the frequency of transition mutations reached now ∼20-30% at the ends of the reads (Figure 3a and 3c) while high numbers of reads were still retained after the harsh filtering, i.e. 43,682 reads for *P. destructans* (3.02% of the genome covered), and 29,647 reads for *P. verrucosus* (3.62% of the genome covered) (Figure 4 a-b-c-d). Nevertheless, the edit distance after PMD filtering in Figure 4a and 4c still demonstrated a relatively high mismatch rate, implying that the filtering procedure did not eliminate potential mis-mapping (i.e., mapping to correct genus but not exactly that species). The phylogenetic relationships of the ancient reads aligned against the two potential fungal candidates were analyzed via an alignment-free approach based on k-mer statistics. Cluster analysis derived from k-mer frequency matrices of fungal species can provide valuable information regarding the taxonomic relationship among the different species (26). Briefly, taxa grouping together are expected to have more similar genetic characteristics than the ones in the other clusters (Figure 5). Although separately in a subcluster, both fungal candidates grouped in the cluster with the other *Pseudogymnoascus* and not with any outgroup species. This further confirms that we robustly identified *Pseudogymnoascus* genus in the Iceman data. However, the exact species is less confident. The two ancient candidates formed a separate subcluster, most likely due to the damage to their genomic data (Figure 5). After the different validation steps, we examined the distribution of the mapped reads of *P. destructans* and *P. verrucosus* along the different body sites of the Iceman. In both cases, the fungal reads were particularly abundant in the stomach, and, to a lesser extent, in the small intestine compartments (Figure 6) (Table 2). Very little to no reads were assigned to the two fungi in the large intestine, muscle and extraction blank (Figure 5) (Table 2).

**Table 1.**
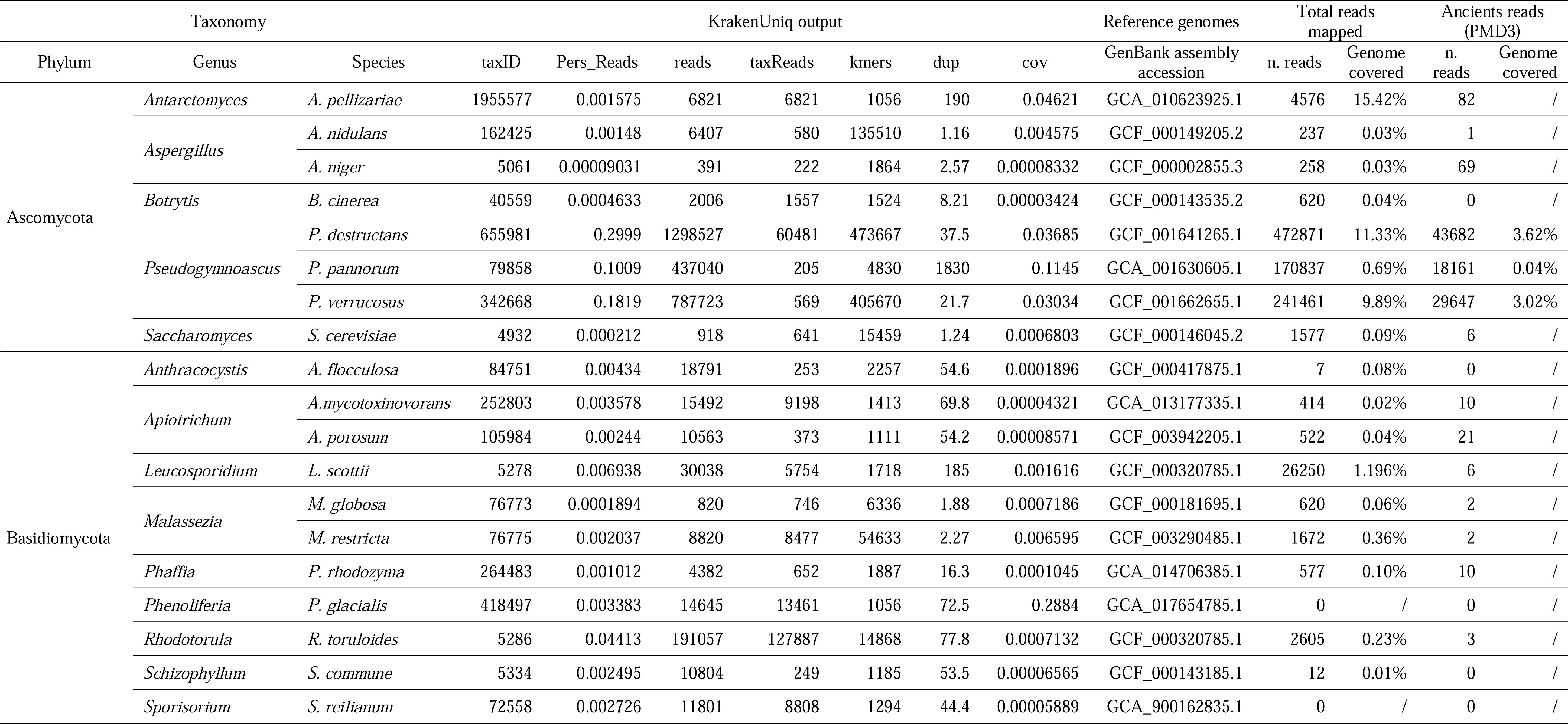
Detection, validation and authentication metrics for fungal species discovered in the Iceman tissues.

**Figure 3.**
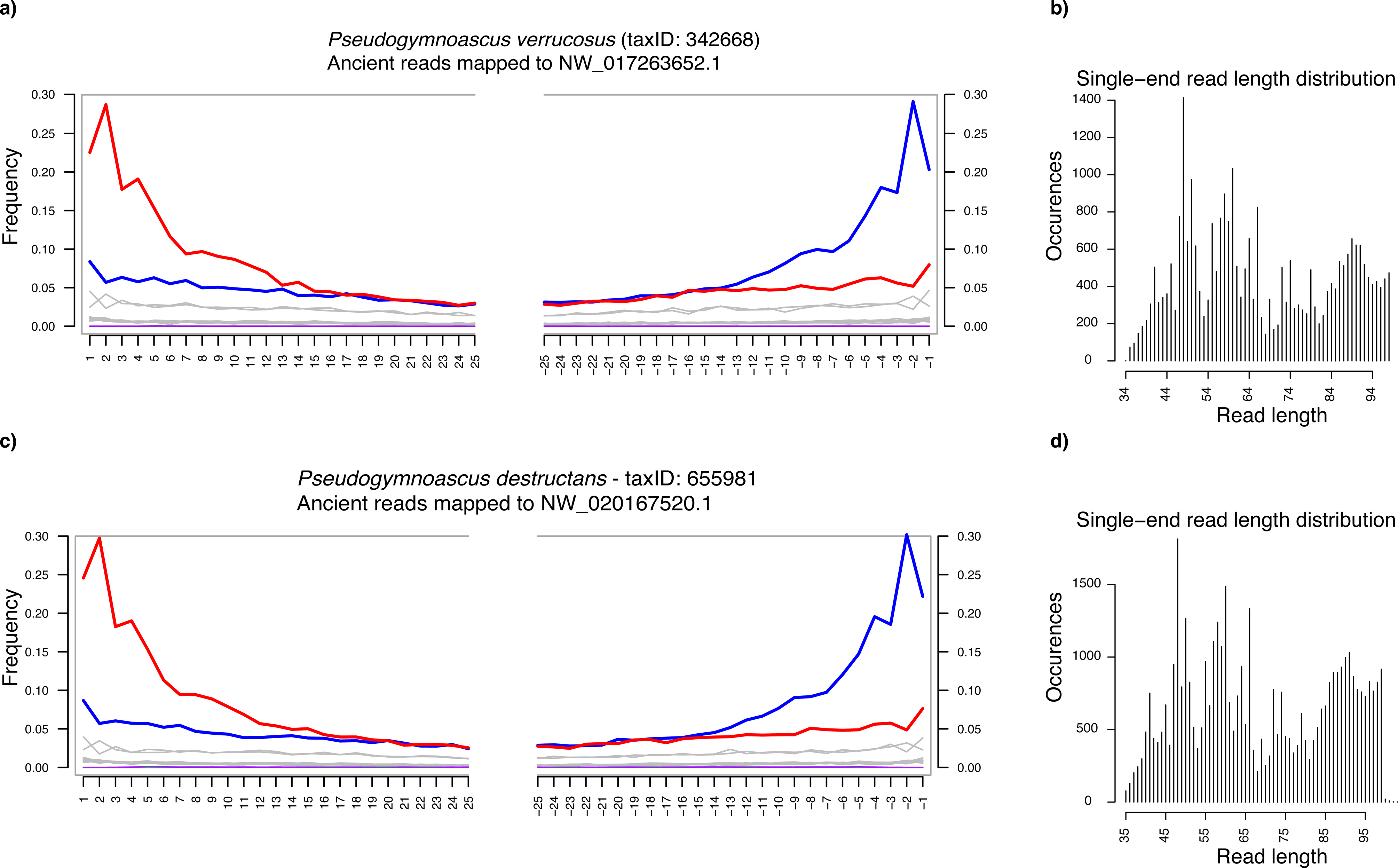
Deamination pattern and read length distribution of *P. verrucosus* (a-b) and *P. destructans* (c-d) after filtering for read length (max. 100 bp long reads were selected) and damage (reads with PMD score > 3 were selected).

**Figure 4.**
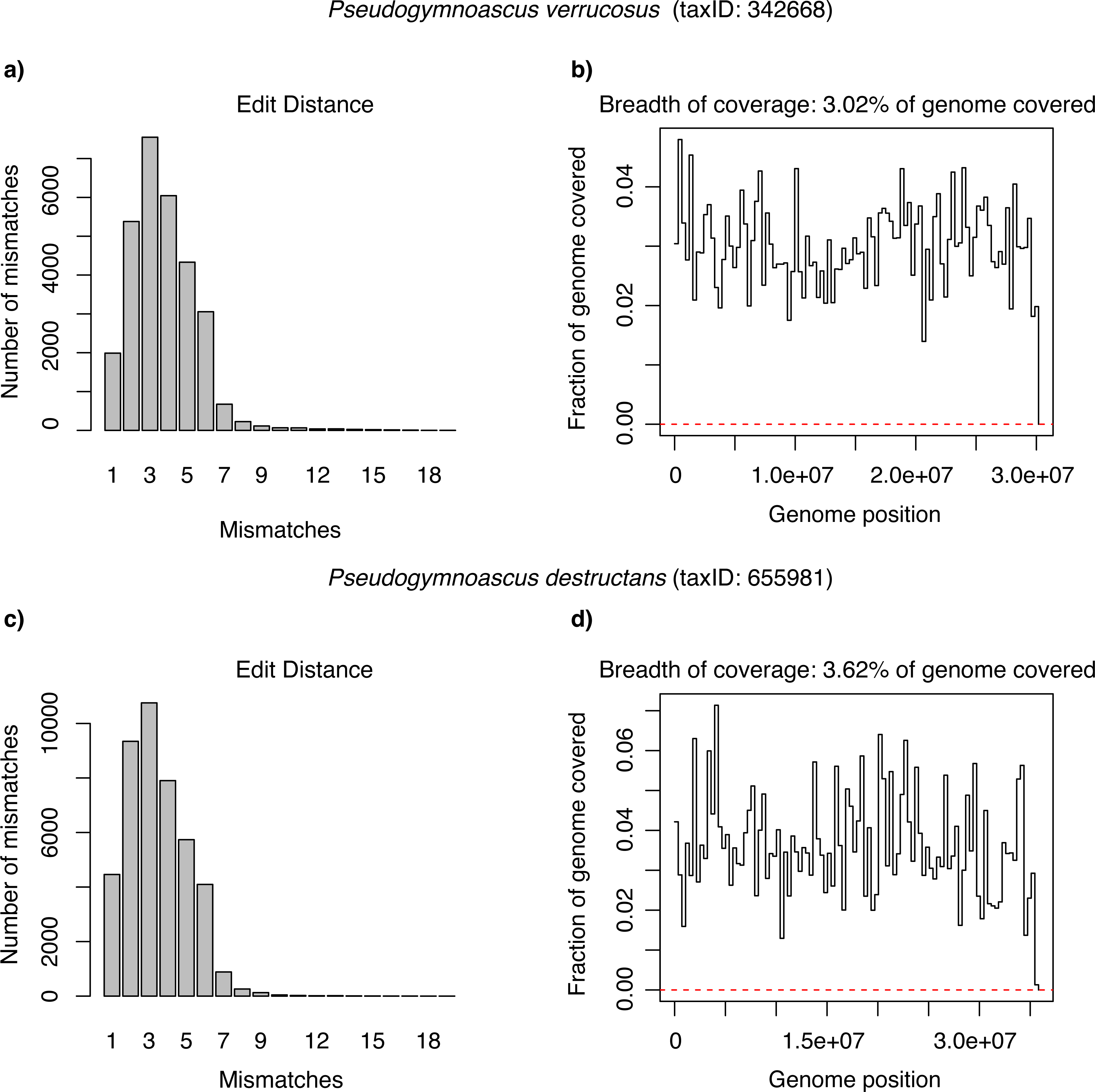
Edit distance and breadth of coverage of *P. verrucosus* (a and b) and *P. destructans ancients* reads (PMD score > 3) (c and d)

**Figure 5.**
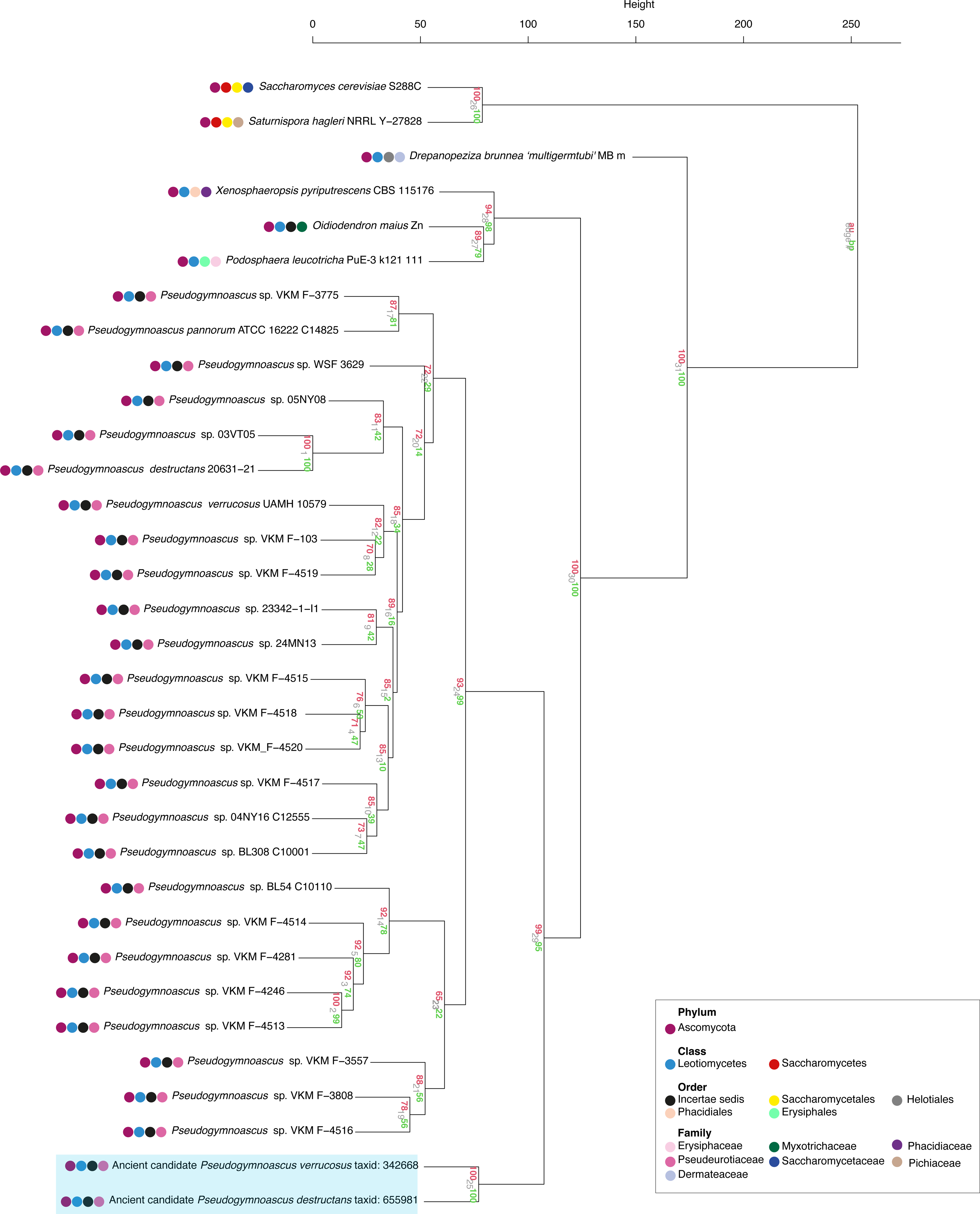
Dendrogram based on Manhattan distance and built using the k-mer method. The dendrogram was constructed using all the genomes of the species belonging to Pseudogymnoascus available in NCBI, along with four outgroup species belonging to the same taxonomic phylum (Ascomicota) and class (Leotiomycetes) as *Pseudogymnoascus*, and two from the same phylum (Ascomicota). The two ancient candidates are highlighted in blue.

**Figure 6.**
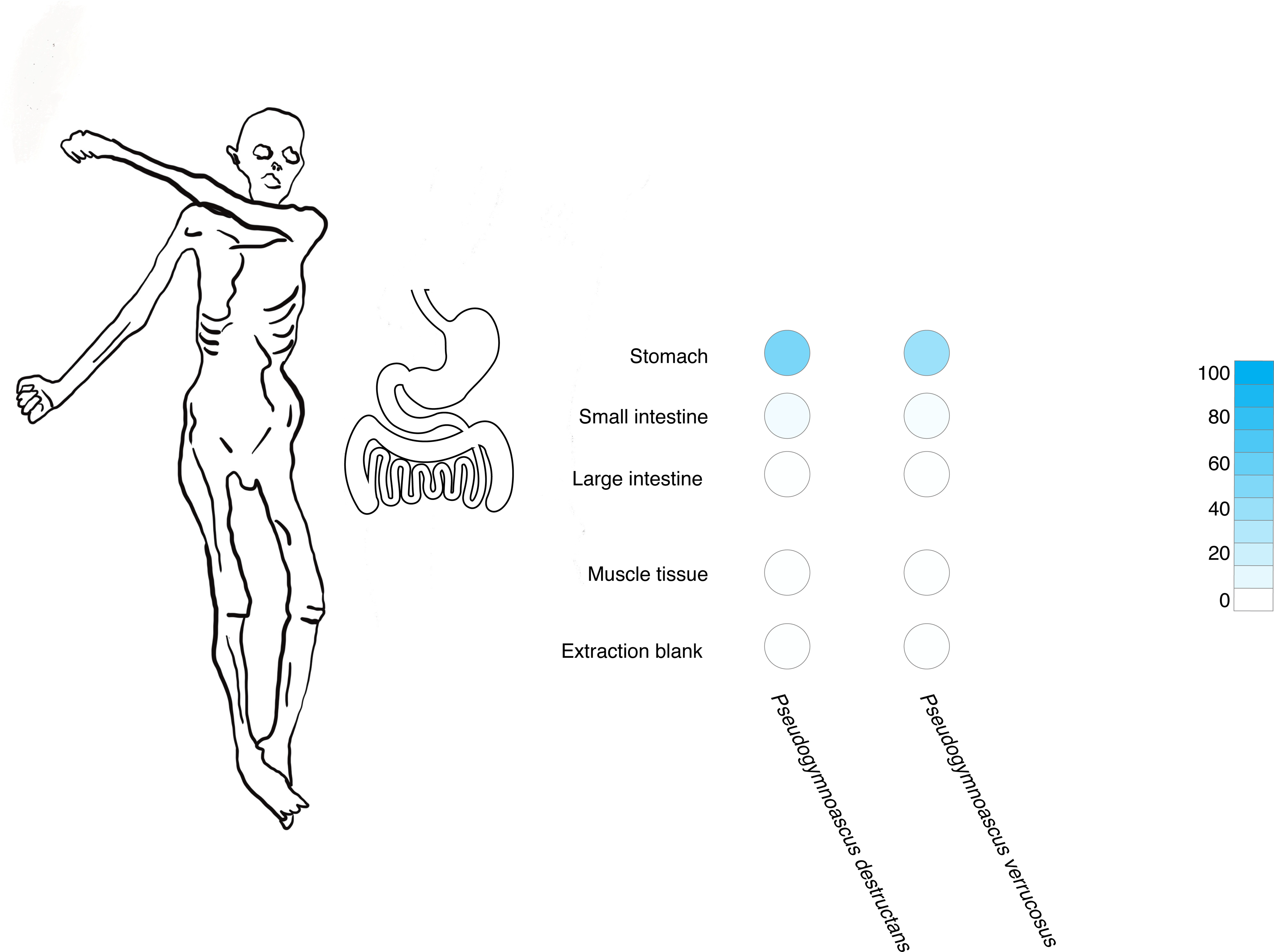
Relative abundance of *P. verrucosus* and *P. destructans* in different Iceman samples.

**Table 2.** Absolute number and frequency of the ancient reads along the different body sites.

|  | *P. verrucosus* (taxID:<br>342668) |  | *P. destructans* (taxID:<br>655981) |  |
| --- | --- | --- | --- | --- |
| Body site | Number<br>of ancient<br>reads | Frequency<br>of ancient<br>reads | Number<br>of ancient<br>reads | Frequency<br>of ancient<br>reads |
| Stomach | 28969 | 3.6 % | 42770 | 5.3 |
| Small intestine | 677 | 38.9% | 910 | 52.3 |
| Large intestine Upper part | > 5 | / | > 5 | / |
| Muscle tissue | > 5 | / | > 5 | / |
| Extraction Blank | > 5 | / | > 5 | / |

The presence of *Pseudogymnoascus* in the gut of the Iceman presents a complex puzzle. *Pseudogymnoascus* encompasses species that are psychrophilic or psychrotolerant saprotrophs, commonly found in diverse environments like soil and associated with plant material (37). Additionally, *P. destructans* can act as opportunistic pathogens, causing infections in the skin and respiratory tract (37, 38). Palmer et al. suggested that *P. destructans* is likely a true fungal pathogen of bats, the causative agent of white-nose syndrome, evolving alongside Eurasian bat species for millions of years (39). Our proposed explanation is that the presence of *Pseudogymnoascus* in the Iceman’s gut may reflect his dietary habits and the broader environmental conditions of his time. It is plausible that the Iceman could have accidentally ingested the fungal species through the diet. After the iceman’s death, the fungus found suitable survival and growth conditions for its proliferation. This hypothesis is supported by the ecology of the *Pseudogymnoascus* species, which has a long-term persistence and can survive and grow at a temperature lower than -20 °C and in harsh environments (40, 38, 41). Moreover, there is evidence that some *Pseudogymnoascus* species, such as *P. destructans* can survive in the gastrointestinal tract (42).

Our research delved into the realm of historical metagenomic samples to unravel the presence of fungal aDNA. Despite the rarity of such findings, we successfully identified the presence of the *Pseudogymnoascus* genus in the Iceman, providing valuable insights into their coexistence with ancient humans. Furthermore, we developed a conservative bioinformatic approach that can potentially be applied to other types of ancient sampling to detect and validate the possible presence of aDNA from filamentous fungi and yeasts.

## Supplementary Information

### Quality control

For the quality analysis of the ancient reads we use bbduk software version xx (cit). Right trimming parameter (ktrim) has been set equal to “r”; k-mer size has been set to 23 and the hamming distance has been set to allow one mismatch (hdist =1). Bbduk was also used to merge the overlapping reads with a minimum length parameter set to 15.

### Alignment

Mapping of trimmed reads to a reference genome was performed using Bowtie2 aligner according to the following command line:

*time bowtie2 --large-index -x ref.fasta --end-to-end --threads 4 --very-sensitive - sample.trimmed.fastq.gz | samtools view -bS -q 1 -h -@ 4 - | samtools sort > sample.trimmed..aligned_to_ref.bam*

### K-mer based profiling with KrakenUniq

Metagenomic samples were taxonomically profiled with KrakenUniq tool (26) using full non-redundant NCBI NT database, and filtered with respect to depth (>200 assigned reads) and breadth of coverage (>1000 unique k-mers) which ensures a good balance between sensitivity and specificity of organism detection (14). The KrakenUniq tool utilizes a Lowest common Ancestor (LCA) algorithm that can deal with DNA reads mapping with the same affinity to multiple organisms.

### Authentication analysis

We computed a number of quality metrics for authenticating the organisms detected in the metagenomic samples. The evenness of coverage was addressed by *samtools depth*, and deamination profile and read length distribution were calculated by *mapDamage* (28). In addition, we computed the edit distance using a custom script, and post-mortem damage (PMD) score distribution via PMDtools (31). The ancient status of detected organisms was addressed, first, by inspecting the incremental pattern of transition mutations frequencies at the terminal ends of the DNA reads, and, second, DNA fragmentation via reads length distribution.

## Tables and Figures

**Figure 1 S.M** Krona charts representation of taxonomic classification of all reads found in the iceman samples

**Figure 2 S.M** Krona charts representation of taxonomic classification of abacterial reads found in the iceman samples

**Figure 3 S.M** Krona charts representation of taxonomic classification of fungal reads found in the iceman samples

**Figure 4 S.M.** Edit distance plots of ancient, aligned reads on *P. verrucosus* (a) and *P. destructans* (b)

**Figure 5 S.M.** Deamination pattern and read length distribution of *P. verrucosus* (a-b) and *P. destructans* (c-d) without PMD filtering.

## References

1. Rasmussen M, Li Y, Lindgreen S, Pedersen JS, Albrechtsen A, Moltke I, et al. Ancient human genome sequence of an extinct Palaeo-Eskimo. Nature. 2010 Feb;463(7282):757–2.62.

2. Van Der Valk T, Pečnerová P, Díez-del-Molino D, Bergström A, Oppenheimer J, Hartmann S, et al. Million-year-old DNA sheds light on the genomic history of mammoths. Nature. 2021 Mar 11;591(7849):265–9.

3. Skoglund P, Ersmark E, Palkopoulou E, Dalén L. Ancient Wolf Genome Reveals an Early Divergence of Domestic Dog Ancestors and Admixture into High-Latitude Breeds. Curr Biol. 2015 Jun;25(11):1515–9.

4. Scott MF, Botigué LR, Brace S, Stevens CJ, Mullin VE, Stevenson A, et al. A 3,000-year-old Egyptian emmer wheat genome reveals dispersal and domestication history. Nat Plants. 2019 Nov 4;5(11):1120–8.

5. Fracasso I, Dinella A, Giammarchi F, Marinchel N, Kołaczek P, Lamentowicz M, et al. Climate and human impacts inferred from a 1500-year multi-proxy record of an alpine peatland in the South-Eastern Alps. Ecol Indic. 2022 Dec;145:109737.

6. Weyrich LS, Duchene S, Soubrier J, Arriola L, Llamas B, Breen J, et al. Neanderthal behaviour, diet, and disease inferred from ancient DNA in dental calculus. Nature. 2017 Apr;544(7650):357–61.

7. Vernot B, Zavala EI, Gómez-Olivencia A, Jacobs Z, Slon V, Mafessoni F, et al. Unearthing Neanderthal population history using nuclear and mitochondrial DNA from cave sediments. Science. 2021 May 7;372(6542):eabf1667.

8. Pedersen MW, De Sanctis B, Saremi NF, Sikora M, Puckett EE, Gu Z, et al. Environmental genomics of Late Pleistocene black bears and giant short-faced bears. Curr Biol. 2021 Jun;31(12):2728–2736.e8.

9. Wang Y, Pedersen MW, Alsos IG, De Sanctis B, Racimo F, Prohaska A, et al. Late Quaternary dynamics of Arctic biota from ancient environmental genomics. Nature. 2021 Dec 2;600(7887):86–92.

10. Rasmussen S, Allentoft ME, Nielsen K, Orlando L, Sikora M, Sjögren KG, et al. Early Divergent Strains of Yersinia pestis in Eurasia 5,000 Years Ago. Cell. 2015 Oct;163(3):571–82.

11. Sarhan MS, Lehmkuhl A, Straub R, Tett A, Wieland G, Francken M, et al. Ancient DNA diffuses from human bones to cave stones. iScience. 2021 Dec;24(12):103397.

12. Maixner F, Krause-Kyora B, Turaev D, Herbig A, Hoopmann MR, Hallows JL, et al. The 5300-year-old *Helicobacter pylori* genome of the Iceman. Science. 2016 Jan 8;351(6269):162–5.

13. Key FM, Posth C, Krause J, Herbig A, Bos KI. Mining Metagenomic Data Sets for Ancient DNA: Recommended Protocols for Authentication. Trends Genet. 2017 Aug;33(8):508–20.

14. Pochon, Z., Bergfeldt, N., Kırdök, E. et al. aMeta: an accurate and memory-efficient ancient metagenomic profiling workflow. Genome Biol 24, 242 (2023). 10.1186/s13059-023-03083-9

15. Mande SS, Mohammed MH, Ghosh TS. Classification of metagenomic sequences: methods and challenges. Brief Bioinform. 2012 Nov 1;13(6):669–81.

17. De Vries RP, Tsang A, Grigoriev IV, editors. Fungal Genomics: Methods and Protocols [Internet]. New York, NY: Springer New York; 2018 [cited 2023 Jun 22]. (Methods in Molecular Biology; vol. 1775). Available from: http://link.springer.com/10.1007/978-1-4939-7804-5

17. Wu B, Hussain M, Zhang W, Stadler M, Liu X, Xiang M. Current insights into fungal species diversity and perspective on naming the environmental DNA sequences of fungi. Mycology. 2019 Jul 3;10(3):127–40.

18. Tedersoo L, Bahram M, Põlme S, Kõljalg U, Yorou NS, Wijesundera R, et al. Global diversity and geography of soil fungi. Science. 2014 Nov 28;346(6213):1256688.

19. Borruso L, Checcucci A, Torti V, Correa F, Sandri C, Luise D, et al. I Like the Way You Eat It: Lemur (Indri indri) Gut Mycobiome and Geophagy. Microb Ecol. 2021 Jul;82(1):215–23.

20. Huseyin CE, O’Toole PW, Cotter PD, Scanlan PD. Forgotten fungi—the gut mycobiome in human health and disease. FEMS Microbiol Rev. 2017 Jul 1;41(4):479– 511.

21. Lavrinienko A, Scholier T, Bates ST, Miller AN, Watts PC. Defining gut mycobiota for wild animals: a need for caution in assigning authentic resident fungal taxa. Anim Microbiome. 2021 Dec;3(1):75.

22. Müller W, Fricke H, Halliday AN, McCulloch MT, Wartho JA. Origin and Migration of the Alpine Iceman. Science. 2003 Oct 31;302(5646):862–6.

23. Mancabelli L, Turroni F, Ferrario C, Duranti S, Van Sinderen D, Ventura M. Ancient bacteria of the Ötzi’s microbiome: a genomic tale from the Copper Age. Microbiome. 2017 Dec;5(1):5.

25. Bushnell B, Rood J, Singer E. BBMerge – Accurate paired shotgun read merging via overlap. Biggs PJ, editor. PLOS ONE. 2017 Oct 26;12(10):e0185056.

25. Andrews S. FastQC: a quality control tool for high throughput sequence data. 2010. 2017;

26. Breitwieser FP, Baker DN, Salzberg SL. KrakenUniq: confident and fast metagenomics classification using unique k-mer counts. Genome Biol. 2018 Nov 16;19(1):198.

27. Langmead B, Salzberg SL. Fast gapped-read alignment with Bowtie 2. Nat Methods. 2012 Apr;9(4):357–9.

28. Jónsson H, Ginolhac A, Schubert M, Johnson PLF, Orlando L. mapDamage2.0: fast approximate Bayesian estimates of ancient DNA damage parameters. Bioinformatics. 2013 Jul;29(13):1682–4.

29. Li H, Handsaker B, Wysoker A, Fennell T, Ruan J, Homer N, et al. The Sequence Alignment/Map format and SAMtools. Bioinformatics. 2009 Aug 15;25(16):2078–9.

30. Altschul SF, Gish W, Miller W, Myers EW, Lipman DJ. Basic local alignment search tool. J Mol Biol. 1990;215(3):403–10.

31. Skoglund P, Northoff BH, Shunkov MV, Derevianko AP, Pääbo S, Krause J, et al. Separating endogenous ancient DNA from modern day contamination in a Siberian Neandertal. Proc Natl Acad Sci. 2014 Feb 11;111(6):2229–34.

32. Ondov BD, Bergman NH, Phillippy AM. Interactive metagenomic visualization in a Web browser. BMC Bioinformatics. 2011 Dec;12(1):385.

33. Marçais G, Kingsford C. A fast, lock-free approach for efficient parallel counting of occurrences of *k* -mers. Bioinformatics. 2011 Mar 15;27(6):764–70.

34. Jari Oksanen, Gavin L. Simpson, F. Guillaume Blanchet, Roeland Kindt, Pierre Legendre, Peter R. Minchin, R.B. O’Hara, Peter Solymos, M. Henry H. Stevens, Eduard Szoecs, Helene Wagner, Matt Barbour, Michael Bedward, Ben Bolker, Daniel Borcard, Gustavo Carvalho, Michael Chirico, Miquel De Caceres, Sebastien Durand, Heloisa Beatriz Antoniazi Evangelista, Rich FitzJohn, Michael Friendly, Brendan Furneaux, Geoffrey Hannigan, Mark O. Hill, Leo Lahti, Dan McGlinn, Marie-Helene Ouellette, Eduardo Ribeiro Cunha, Tyler Smith, Adrian Stier, Cajo J.F. Ter Braak, James Weedon. vegan: Community Ecology Packag. Ordination methods, diversity analysis and other functions for community and vegetation ecologists.

35. Suzuki R, Shimodaira H. Pvclust: an R package for assessing the uncertainty in hierarchical clustering. Bioinformatics. 2006 Jun 15;22(12):1540–2.

36. Briggs AW, Stenzel U, Johnson PLF, Green RE, Kelso J, Prüfer K, et al. Patterns of damage in genomic DNA sequences from a Neandertal. Proc Natl Acad Sci. 2007 Sep 11;104(37):14616–21.

37. Quandt CA, Haelewaters D. Phylogenetic Advances in Leotiomycetes, an Understudied Clade of Taxonomically and Ecologically Diverse Fungi. In: Zaragoza Ó, Casadevall A, editors. Encyclopedia of Mycology [Internet]. Oxford: Elsevier; 2021. p. 284–94. Available from: https://www.sciencedirect.com/science/article/pii/B9780128199909000524

38. Hayes MA. The Geomyces Fungi: Ecology and Distribution. BioScience. 2012 Sep;62(9):819–23.

39. Palmer JM, Drees KP, Foster JT, Lindner DL. Extreme sensitivity to ultraviolet light in the fungal pathogen causing white-nose syndrome of bats. Nat Commun. 2018 Jan 2;9(1):35.

41. Veselská et al. - 2020 - Comparative eco-physiology revealed extensive enzy.pdf.

41. Hoyt JR, Langwig KE, Okoniewski J, Frick WF, Stone WB, Kilpatrick AM. Long-Term Persistence of *Pseudogymnoascus destructans*, the Causative Agent of White-Nose Syndrome, in the Absence of Bats. EcoHealth. 2015 Jun;12(2):330–3.

42. Ballmann AE, Torkelson MR, Bohuski EA, Russell RE, Blehert DS. Dispersal hazards of *Pseudogymnoascus destructans* by bats and human activity at hibernacula in summer. J Wildl Dis. 2017 Oct 1;53(4):725.

