## Supplementary figures and images for "Unravelling the ancient fungal DNA from the Iceman’s gut"

### Figure 1 Supplementary

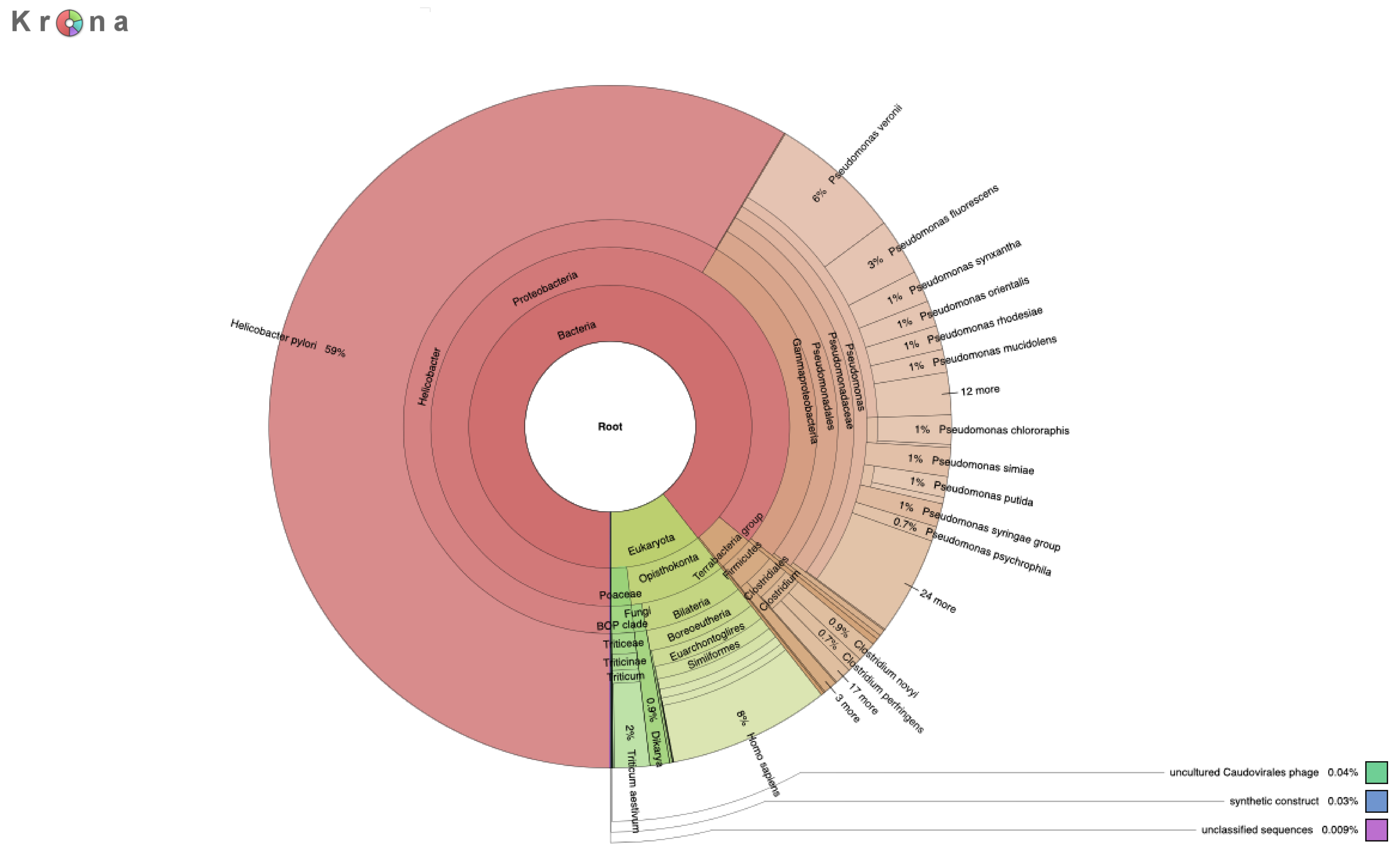

### Figure 2 Supplementary

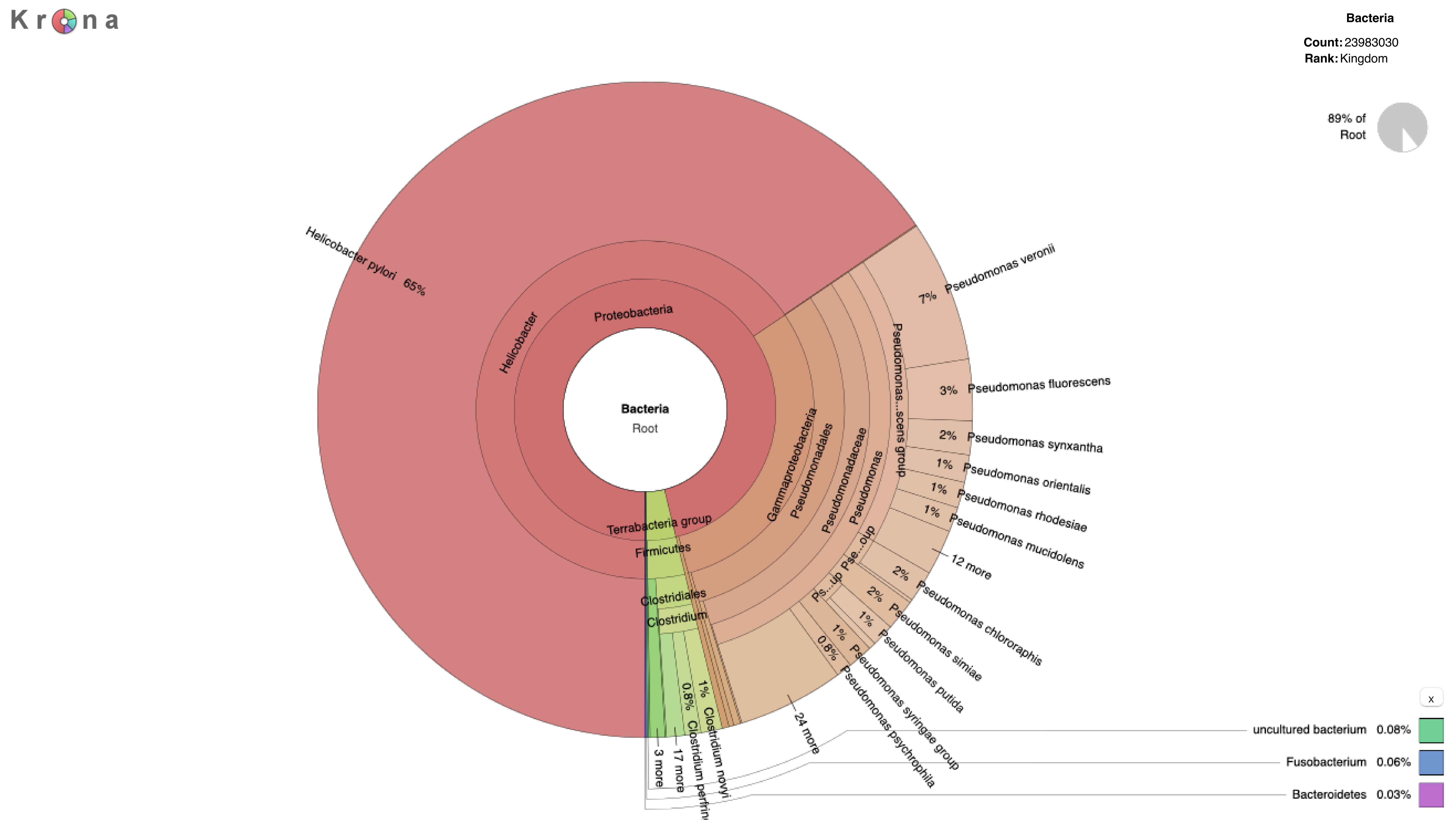

### Figure 3 Supplementary

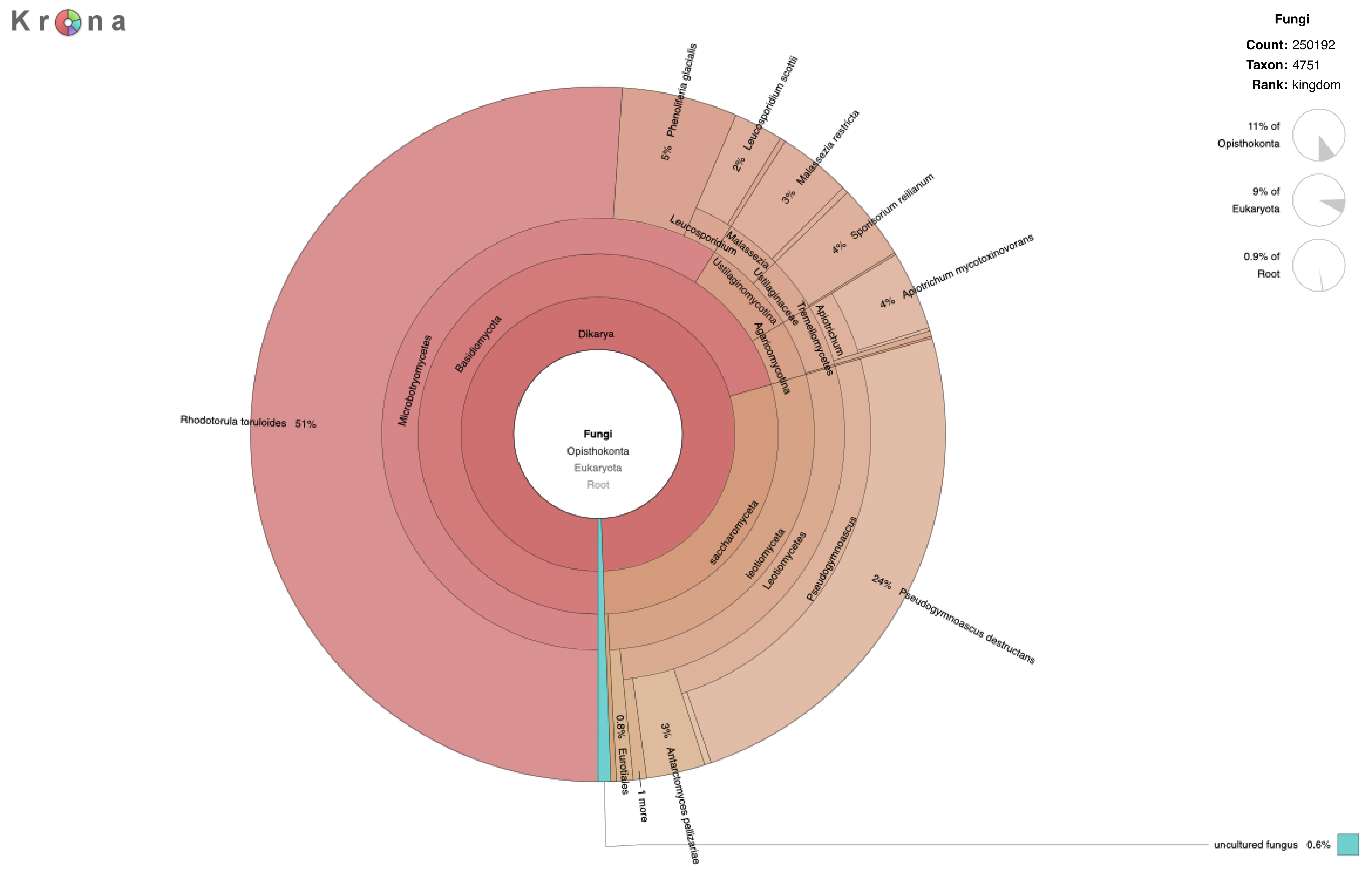

### Figure 4a-b Supplementary

**a)**

*Pseudogymnoascus verrucosus* (taxID: 342668)

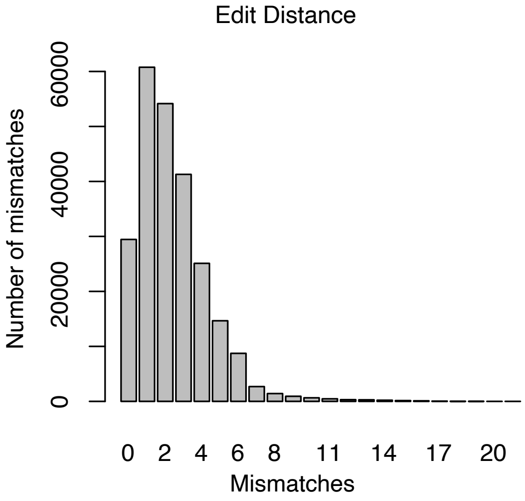


**b)**

*Pseudogymnoascus destructans* (taxID: 655981)

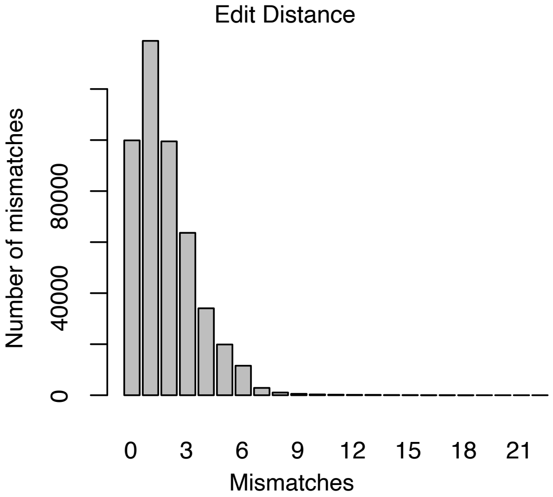

### Figure 5a-b-c-d Supplementary

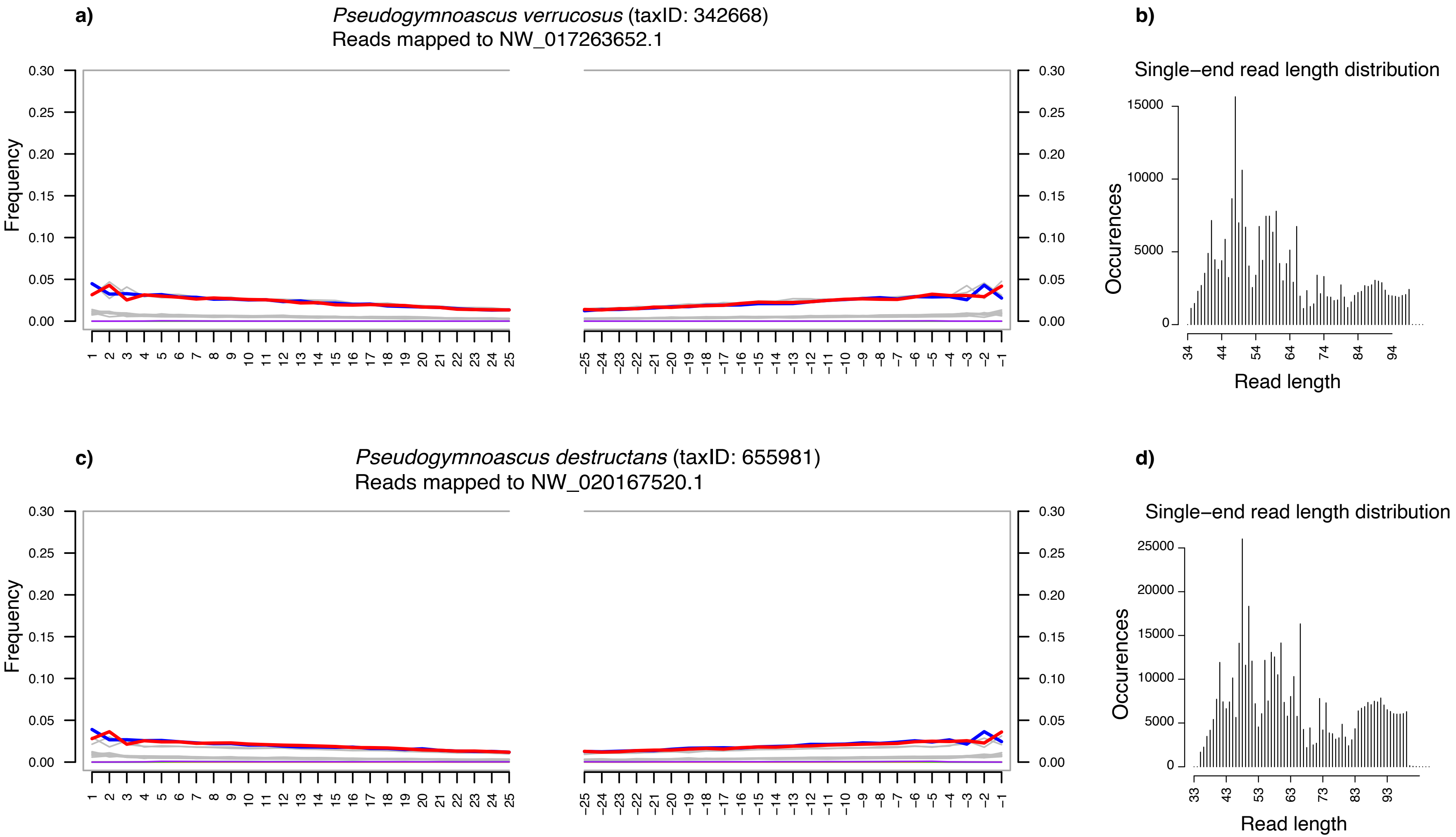
