## Supplementary material for "Unravelling the ancient fungal DNA from the Iceman’s gut": Table 1 Supplementary

| taxName (Kingdom) | taxName (Species) | taxID | Pers_Reads | reads | taxReads | kmers | dup | cov |
|---|---|---|---|---|---|---|---|---|
| Animalia | Bos mutus | 72004 | 0.02389 | 103435 | 15676 | 837327 | 1.18 | 0.001219 |
| Animalia | Bos taurus | 9913 | 0.001606 | 6952 | 1621 | 57759 | 1.21 | 0.001493 |
| Animalia | Bubalus bubalis | 89462 | 0.0004192 | 1815 | 525 | 17995 | 1.16 | 0.0005988 |
| Animalia | Callithrix jacchus | 9483 | 0.0002511 | 1087 | 314 | 5641 | 1.43 | 6.78E-05 |
| Animalia | Capra hircus | 9925 | 0.003139 | 13590 | 3345 | 105669 | 2.01 | 0.006128 |
| Animalia | Capra ibex | 72542 | 0.002183 | 9451 | 1625 | 1788 | 103 | 0.5598 |
| Animalia | Cervus elaphus | 9860 | 0.002133 | 9233 | 2041 | 5370 | 22 | 0.07398 |
| Animalia | Cervus nippon | 9863 | 0.0009796 | 4241 | 1002 | 1048 | 43.2 | 0.007449 |
| Animalia | Gorilla gorilla | 9593 | 0.0003545 | 1535 | 302 | 9184 | 1.48 | 0.0004862 |
| Animalia | Homo sapiens | 9606 | 0.4736 | 2050251 | 2049978 | 45042725 | 1.41 | 0.03044 |
| Animalia | Macaca mulatta | 9544 | 0.0008946 | 3873 | 779 | 22256 | 1.37 | 0.0001765 |
| Animalia | Muntiacus vaginalis | 9887 | 0.0004938 | 2138 | 258 | 2311 | 5.72 | 0.003767 |
| Animalia | Mus musculus | 10090 | 0.003105 | 13445 | 13442 | 4287 | 42.6 | 2.19E-06 |
| Animalia | Ovis aries | 9940 | 0.001024 | 4435 | 994 | 50576 | 1.23 | 0.002629 |
| Animalia | Pan paniscus | 9597 | 0.0002023 | 876 | 336 | 5252 | 1.73 | 0.0007113 |
| Animalia | Pan troglodytes | 9598 | 0.003462 | 14990 | 5732 | 105050 | 1.5 | 0.0009041 |
| Animalia | Pongo abelii | 9601 | 0.00104 | 4502 | 1122 | 27391 | 1.44 | 0.0003081 |
| Animalia | Sus scrofa | 9823 | 0.0006144 | 2660 | 706 | 29152 | 1.33 | 0.0001003 |
| Animalia | Caenorhabditis elegans | 6239 | 0.000137 | 593 | 466 | 38576 | 1.11 | 0.0004441 |
| Animalia | Spirometra erinaceieuropaei | 99802 | 0.001059 | 4585 | 280 | 12339 | 5.33 | 1.43E-05 |
| Bacteria | Arthrobacter sp. | 1667 | 9.03E-05 | 391 | 322 | 5016 | 1.91 | 0.007851 |
| Bacteria | Corynebacterium xerosis | 1725 | 0.0002682 | 1161 | 432 | 13310 | 1.02 | 0.003326 |
| Bacteria | Cutibacterium acnes | 1747 | 0.003481 | 15072 | 14782 | 601625 | 1.73 | 0.1412 |
| Bacteria | Kocuria rhizophila | 72000 | 0.000252 | 1091 | 1090 | 21148 | 1.46 | 0.04028 |
| Bacteria | Pseudopropionibacterium propionicum | 1750 | 0.0001261 | 546 | 380 | 10709 | 1.12 | 0.001981 |
| Bacteria | Rhodococcus erythropolis | 1833 | 0.000131 | 567 | 477 | 2995 | 1.82 | 0.0005314 |
| Bacteria | Bradyrhizobium diazoefficiens | 1355477 | 0.0002901 | 1256 | 1203 | 1042 | 12.1 | 0.0001456 |
| Bacteria | Bradyrhizobium symbiodeficiens | 1404367 | 6.42E-05 | 278 | 208 | 1282 | 2.54 | 0.0001583 |
| Bacteria | Caulobacter segnis | 88688 | 0.000155 | 671 | 670 | 3216 | 2.87 | 0.0007915 |
| Bacteria | Caulobacter vibrioides | 155892 | 0.0001631 | 706 | 256 | 3688 | 4.06 | 0.000462 |
| Bacteria | uncultured bacterium | 77133 | 0.01532 | 66306 | 18478 | 86648 | 7.33 | 0.0003989 |
| Bacteria | Flavobacterium anhuiense | 459526 | 0.0001446 | 626 | 278 | 3251 | 1.54 | 0.000534 |
| Bacteria | Flavobacterium johnsoniae | 986 | 0.0003645 | 1578 | 687 | 13102 | 1.47 | 0.001786 |
| Bacteria | Porphyromonas gingivalis | 837 | 7.55E-05 | 327 | 242 | 10357 | 1.11 | 0.001767 |
| Bacteria | Prevotella denticola | 28129 | 0.0001714 | 742 | 351 | 17421 | 1.16 | 0.004637 |
| Bacteria | Prevotella intermedia | 28131 | 0.0003481 | 1507 | 965 | 32683 | 1.62 | 0.003104 |
| Bacteria | Prevotella melaninogenica | 28132 | 0.0009786 | 4237 | 1289 | 60347 | 1.27 | 0.009168 |

| | | | | | | | | |
|---|---|---|---|---|---|---|---|---|
| Bacteria | Tannerella sp. oral taxon HOT-286 | 712710 | 0.000124 | 537 | 533 | 10993 | 1.16 | 0.003896 |
| Bacteria | Gemella morbillorum | 29391 | 0.0005562 | 2408 | 1283 | 55874 | 1.29 | 0.02919 |
| Bacteria | Alkalihalobacillus clausii | 79880 | 0.0001266 | 548 | 437 | 1459 | 1.79 | 0.0001902 |
| Bacteria | Brochothrix thermosphacta | 2756 | 9.54E-05 | 413 | 370 | 7955 | 1.53 | 0.00224 |
| Bacteria | Carnobacterium maltaromaticum | 2751 | 0.002399 | 10388 | 7439 | 373528 | 1.29 | 0.06895 |
| Bacteria | Clostridioides difficile | 1496 | 0.001608 | 6960 | 6674 | 6472 | 10.3 | 0.0005715 |
| Bacteria | Clostridium acetobutylicum | 1488 | 0.002844 | 12314 | 12047 | 4069 | 25.8 | 0.001101 |
| Bacteria | Clostridium argentinense | 29341 | 0.005959 | 25798 | 238 | 7092 | 31.6 | 0.001733 |
| Bacteria | Clostridium baratii | 1561 | 0.003788 | 16400 | 7384 | 27141 | 5.65 | 0.006082 |
| Bacteria | Clostridium beijerinckii | 1520 | 0.01261 | 54593 | 7546 | 26483 | 17.8 | 0.00202 |
| Bacteria | Clostridium botulinum | 1491 | 0.0288 | 124672 | 69866 | 52477 | 23.7 | 0.001757 |
| Bacteria | Clostridium butyricum | 1492 | 0.003486 | 15092 | 12183 | 27914 | 4.91 | 0.003568 |
| Bacteria | Clostridium cellulovorans | 1493 | 0.002551 | 11045 | 1125 | 2880 | 28.2 | 0.0006027 |
| Bacteria | Clostridium chauvoei | 46867 | 0.001896 | 8210 | 8174 | 3436 | 18.6 | 0.00148 |
| Bacteria | Clostridium estertheticum | 238834 | 1.34 | 5803423 | 515 | 1312068 | 81.4 | 0.3102 |
| Bacteria | Clostridium kluyveri | 1534 | 0.004622 | 20013 | 10176 | 5938 | 23.2 | 0.0009385 |
| Bacteria | Clostridium novyi | 1542 | 0.5127 | 2219890 | 252173 | 1855997 | 37.9 | 0.5062 |
| Bacteria | Clostridium pasteurianum | 1501 | 0.0163 | 70581 | 22306 | 16431 | 35.9 | 0.002054 |
| Bacteria | Clostridium perfringens | 1502 | 0.07011 | 303524 | 193388 | 3218107 | 2.75 | 0.2186 |
| Bacteria | Clostridium saccharobutylicum | 169679 | 0.004158 | 18003 | 17136 | 6915 | 21.8 | 0.001466 |
| Bacteria | Clostridium saccharoperbutylacetonicum | 36745 | 0.004955 | 21454 | 12397 | 6975 | 23.5 | 0.0009043 |
| Bacteria | Clostridium septicum | 1504 | 0.002448 | 10597 | 10431 | 3528 | 24 | 0.001245 |
| Bacteria | Clostridium tetani | 1513 | 0.003996 | 17302 | 11001 | 8115 | 17.7 | 0.002025 |
| Bacteria | Clostridium tyrobutyricum | 1519 | 0.002826 | 12234 | 10789 | 3503 | 25.2 | 0.001061 |
| Bacteria | Enterococcus faecalis | 1351 | 0.000291 | 1260 | 597 | 19867 | 1.48 | 0.001816 |
| Bacteria | Finegoldia magna | 1260 | 0.0001453 | 629 | 346 | 2931 | 4.97 | 0.00115 |
| Bacteria | Lactobacillus curvatus | 28038 | 0.007297 | 31593 | 22237 | 655647 | 1.66 | 0.1452 |
| Bacteria | Lactobacillus mucosae | 97478 | 0.000131 | 567 | 314 | 6877 | 1.17 | 0.002151 |
| Bacteria | Lactobacillus plantarum | 1590 | 0.0001282 | 555 | 218 | 2832 | 2.72 | 0.000206 |
| Bacteria | Lactobacillus sakei | 1599 | 0.03962 | 171520 | 77183 | 1950679 | 3.13 | 0.3298 |
| Bacteria | Lactococcus lactis | 1358 | 0.0002749 | 1190 | 291 | 2681 | 3.2 | 0.0002038 |
| Bacteria | Leuconostoc mesenteroides | 1245 | 0.0001825 | 790 | 489 | 12757 | 1.51 | 0.002469 |
| Bacteria | Listeria monocytogenes | 1639 | 0.0001684 | 729 | 435 | 2706 | 3.64 | 0.0001939 |

| | | | | | | | |
|---|---|---|---|---|---|---|---|
| Bacteria | Paeniclostridium sordellii | 1505 | 0.002722 | 11785 | 8723 | 73848 | 1.83 | 0.01567 |
| Bacteria | Paraclostridium bifermentans | 1490 | 0.001527 | 6611 | 213 | 14148 | 3.7 | 0.004901 |
| Bacteria | Parvimonas micra | 33033 | 0.001317 | 5703 | 3183 | 153606 | 1.35 | 0.07058 |
| Bacteria | Roseburia intestinalis | 166486 | 8.36E-05 | 362 | 226 | 3769 | 1.22 | 0.0007068 |
| Bacteria | Staphylococcus aureus | 1280 | 0.0002157 | 934 | 408 | 2355 | 5.12 | 0.0001584 |
| Bacteria | Staphylococcus capitis | 29388 | 0.0001901 | 823 | 369 | 15858 | 1.54 | 0.003979 |
| Bacteria | Staphylococcus cohnii | 29382 | 7.02E-05 | 304 | 296 | 3120 | 1.6 | 0.0006328 |
| Bacteria | Staphylococcus epidermidis | 1282 | 0.0002836 | 1228 | 1112 | 26210 | 1.44 | 0.003705 |
| Bacteria | Staphylococcus haemolyticus | 1283 | 8.45E-05 | 366 | 278 | 4675 | 1.76 | 0.001264 |
| Bacteria | Staphylococcus hominis | 1290 | 0.0001575 | 682 | 265 | 9282 | 1.57 | 0.001959 |
| Bacteria | Staphylococcus warneri | 1292 | 5.61E-05 | 243 | 202 | 1501 | 1.77 | 0.0006347 |
| Bacteria | Streptococcus anginosus | 1328 | 0.0003677 | 1592 | 1064 | 26990 | 1.27 | 0.01107 |
| Bacteria | Streptococcus cristatus | 45634 | 0.0003626 | 1570 | 752 | 18740 | 1.26 | 0.00463 |
| Bacteria | Streptococcus gordonii | 1302 | 0.006942 | 30055 | 22375 | 504538 | 1.46 | 0.08421 |
| Bacteria | Streptococcus intermedius | 1338 | 0.004376 | 18944 | 16194 | 342439 | 1.61 | 0.1306 |
| Bacteria | Streptococcus mitis | 28037 | 0.000701 | 3035 | 883 | 32992 | 1.26 | 0.0063 |
| Bacteria | Streptococcus oralis | 1303 | 0.002832 | 12263 | 2120 | 126933 | 1.4 | 0.02448 |
| Bacteria | Streptococcus parasanguinis | 1318 | 0.000197 | 853 | 260 | 9096 | 1.33 | 0.00427 |
| Bacteria | Streptococcus pneumoniae | 1313 | 0.0005888 | 2549 | 1558 | 28264 | 1.48 | 0.003902 |
| Bacteria | Streptococcus porcinus | 1340 | 8.87E-05 | 384 | 300 | 1548 | 3.33 | 0.0005033 |
| Bacteria | Streptococcus pyogenes | 1314 | 0.0001215 | 526 | 460 | 1363 | 3.48 | 0.0002268 |
| Bacteria | Streptococcus sanguinis | 1305 | 0.001573 | 6810 | 2568 | 101512 | 1.25 | 0.01097 |
| Bacteria | Streptococcus suis | 1307 | 0.0001626 | 704 | 478 | 1277 | 3.34 | 8.42E-05 |
| Bacteria | Streptococcus thermophilus | 1308 | 7.99E-05 | 346 | 312 | 2190 | 2.23 | 0.0004658 |
| Bacteria | uncultured Clostridium sp. | 59620 | 0.001891 | 8185 | 391 | 1753 | 58.5 | 0.005879 |
| Bacteria | Veillonella parvula | 29466 | 0.002622 | 11354 | 7681 | 188781 | 1.31 | 0.04887 |
| Bacteria | Fusobacterium hwasookii | 1583098 | 0.0009689 | 4195 | 2290 | 58227 | 1.4 | 0.03166 |
| Bacteria | Fusobacterium nucleatum | 851 | 0.004458 | 19299 | 6245 | 254482 | 1.46 | 0.02573 |
| Bacteria | Fusobacterium pseudoperiodonticum | 2663009 | 0.002056 | 8902 | 5230 | 123382 | 1.44 | 0.02828 |
| Bacteria | Fusobacterium ulcerans | 861 | 0.0007553 | 3270 | 1339 | 1047 | 17.4 | 0.0002106 |
| Bacteria | Pseudomonas amygdali | 47877 | 0.005674 | 24566 | 7033 | 4057 | 36.7 | 0.001307 |
| Bacteria | Pseudomonas stutzeri | 316 | 0.07557 | 327183 | 127704 | 73429 | 30 | 0.001736 |

| | | | | | | | | |
|---|---|---|---|---|---|---|---|---|
| Bacteria | Pseudomonas syringae | 317 | 0.131 | 566994 | 112098 | 64662 | 64.4 | 0.002517 |
| Bacteria | Achromobacter denitrificans | 32002 | 0.0001335 | 578 | 493 | 2629 | 4.71 | 0.0002848 |
| Bacteria | Achromobacter insolitus | 217204 | 0.0003088 | 1337 | 1161 | 1334 | 10.6 | 0.0001619 |
| Bacteria | Achromobacter spanius | 217203 | 0.001849 | 8007 | 1360 | 5968 | 7.95 | 0.0004006 |
| Bacteria | Achromobacter xylosoxidans | 85698 | 0.000929 | 4022 | 1231 | 6832 | 7.81 | 0.0002596 |
| Bacteria | Acidovorax carolinensis | 553814 | 0.0003506 | 1518 | 1137 | 5857 | 4.11 | 0.0009392 |
| Bacteria | Acinetobacter baumannii | 470 | 0.01363 | 59010 | 2182 | 422391 | 3.42 | 0.02196 |
| Bacteria | Acinetobacter haemolyticus | 29430 | 0.001588 | 6876 | 2265 | 46909 | 2.61 | 0.006488 |
| Bacteria | Acinetobacter indicus | 756892 | 0.0009066 | 3925 | 1728 | 27481 | 2.47 | 0.003062 |
| Bacteria | Acinetobacter johnsonii | 40214 | 0.02982 | 129102 | 56154 | 1313849 | 2.25 | 0.2222 |
| Bacteria | Acinetobacter junii | 40215 | 0.004177 | 18084 | 10061 | 365734 | 1.64 | 0.08415 |
| Bacteria | Acinetobacter lwoffii | 28090 | 0.001666 | 7213 | 1894 | 91035 | 2.47 | 0.02615 |
| Bacteria | Acinetobacter nosocomialis | 106654 | 0.0003903 | 1690 | 1026 | 6849 | 2.09 | 0.0009016 |
| Bacteria | Acinetobacter pittii | 48296 | 0.0004428 | 1917 | 501 | 16144 | 2.1 | 0.001304 |
| Bacteria | Acinetobacter radioresistens | 40216 | 0.0001871 | 810 | 290 | 7109 | 2.49 | 0.001435 |
| Bacteria | Acinetobacter schindleri | 108981 | 0.0004645 | 2011 | 397 | 17248 | 2 | 0.00265 |
| Bacteria | Acinetobacter wuhouensis | 1879050 | 0.0003402 | 1473 | 411 | 9542 | 2.23 | 0.001834 |
| Bacteria | Aeromonas hydrophila | 644 | 0.0005213 | 2257 | 549 | 1088 | 16.3 | 5.69E-05 |
| Bacteria | Aeromonas salmonicida | 645 | 0.0004107 | 1778 | 1563 | 1779 | 7.96 | 0.0001843 |
| Bacteria | Aeromonas veronii | 654 | 0.001009 | 4370 | 801 | 2478 | 20.8 | 8.26E-05 |
| Bacteria | Aggregatibacter actinomycetemcomitan s | 714 | 0.00068 | 2944 | 2899 | 2102 | 4.87 | 0.0004858 |
| Bacteria | Aggregatibacter aphrophilus | 732 | 0.0008458 | 3662 | 1897 | 94627 | 1.15 | 0.02214 |
| Bacteria | Agrobacterium tumefaciens | 358 | 0.0007922 | 3430 | 1682 | 12423 | 2.79 | 0.0003332 |
| Bacteria | Alcaligenes faecalis | 511 | 0.0006922 | 2997 | 2035 | 3769 | 8.56 | 0.0001989 |
| Bacteria | Alicycliphilus denitrificans | 179636 | 0.0007546 | 3267 | 2120 | 15985 | 1.77 | 0.003142 |
| Bacteria | Azoarcus communis | 41977 | 0.0006352 | 2750 | 373 | 5620 | 5.36 | 0.0007228 |
| Bacteria | Azoarcus olearius | 418699 | 0.0004144 | 1794 | 1406 | 1023 | 8.85 | 0.0002118 |
| Bacteria | Azotobacter chroococcum | 353 | 0.002234 | 9672 | 5226 | 2433 | 44.2 | 0.000434 |
| Bacteria | Azotobacter vinelandii | 354 | 0.00235 | 10174 | 10133 | 2230 | 41.5 | 0.000525 |
| Bacteria | Bordetella hinzii | 103855 | 0.0001072 | 464 | 337 | 1420 | 6.31 | 0.0002382 |
| Bacteria | Buchnera aphidicola | 9 | 0.0006638 | 2874 | 855 | 1158 | 25.6 | 5.94E-05 |
| Bacteria | Burkholderia cenocepacia | 95486 | 0.0007052 | 3053 | 223 | 1213 | 18.7 | 5.18E-05 |
| Bacteria | Burkholderia gladioli | 28095 | 0.001159 | 5018 | 3877 | 1852 | 17 | 0.0001207 |

| | | | | | | | | |
|---|---|---|---|---|---|---|---|---|
| Bacteria | Burkholderia multivorans | 87883 | 0.0003231 | 1399 | 954 | 1318 | 7.01 | 7.77E-05 |
| Bacteria | Cedecea neteri | 158822 | 0.0007509 | 3251 | 1799 | 2327 | 12.3 | 0.0001704 |
| Bacteria | Chryseobacterium bernardetii | 1241978 | 0.0001178 | 510 | 265 | 5067 | 1.4 | 0.0009924 |
| Bacteria | Chryseobacterium carnipullorum | 1124835 | 0.0002097 | 908 | 906 | 10083 | 1.31 | 0.002193 |
| Bacteria | Chryseobacterium gallinarum | 1324352 | 8.92E-05 | 386 | 264 | 2068 | 1.4 | 0.0003923 |
| Bacteria | Chryseobacterium indologenes | 253 | 0.0004825 | 2089 | 1548 | 46940 | 1.4 | 0.003345 |
| Bacteria | Chryseobacterium lactis | 1241981 | 7.97E-05 | 345 | 344 | 1901 | 1.48 | 0.000406 |
| Bacteria | Chryseobacterium nakagawai | 1241982 | 4.69E-05 | 203 | 203 | 1341 | 1.38 | 0.0004166 |
| Bacteria | Chryseobacterium shandongense | 1493872 | 8.57E-05 | 371 | 282 | 1953 | 2.47 | 0.0003928 |
| Bacteria | Citrobacter freundii | 546 | 0.0006585 | 2851 | 790 | 1724 | 8.71 | 6.72E-05 |
| Bacteria | Collimonas arenae | 279058 | 0.001065 | 4609 | 2742 | 6006 | 9.27 | 0.000627 |
| Bacteria | Collimonas fungivorans | 158899 | 0.0007507 | 3250 | 487 | 7894 | 5.38 | 0.0009228 |
| Bacteria | Collimonas pratensis | 279113 | 0.0005114 | 2214 | 1502 | 5017 | 5.82 | 0.0007276 |
| Bacteria | Comamonas testosteroni | 285 | 0.0003028 | 1311 | 512 | 7158 | 3.69 | 0.0005629 |
| Bacteria | Comamonas thiooxydans | 363952 | 0.0004056 | 1756 | 1435 | 7506 | 2.34 | 0.001168 |
| Bacteria | Cupriavidus gilardii | 82541 | 0.001001 | 4334 | 3190 | 1699 | 12.9 | 0.0002913 |
| Bacteria | Cupriavidus metallidurans | 119219 | 0.0006895 | 2985 | 2360 | 39679 | 1.33 | 0.003943 |
| Bacteria | Cupriavidus taiwanensis | 164546 | 0.000534 | 2312 | 689 | 1991 | 10.9 | 0.0001251 |
| Bacteria | Delftia tsuruhatensis | 180282 | 0.0001102 | 477 | 300 | 6227 | 1.69 | 0.001353 |
| Bacteria | Dickeya zeae | 204042 | 0.0004479 | 1939 | 1143 | 1670 | 8.63 | 0.0001309 |
| Bacteria | Eikenella corrodens | 539 | 0.0002617 | 1133 | 269 | 13066 | 1.35 | 0.003868 |
| Bacteria | Enterobacter cloacae | 550 | 0.001898 | 8216 | 1529 | 3200 | 16.8 | 0.0001039 |
| Bacteria | Escherichia coli | 562 | 0.001827 | 7911 | 5407 | 7480 | 7.17 | 9.12E-05 |
| Bacteria | Haemophilus haemolyticus | 726 | 0.000477 | 2065 | 437 | 29664 | 1.41 | 0.009179 |
| Bacteria | Haemophilus influenzae | 727 | 0.001509 | 6533 | 4546 | 108100 | 1.42 | 0.01288 |
| Bacteria | Haemophilus parainfluenzae | 729 | 0.001904 | 8245 | 1775 | 144054 | 1.39 | 0.02592 |
| Bacteria | Helicobacter acinonychis | 212 | 0.001811 | 7842 | 7605 | 6949 | 45.9 | 0.007105 |
| Bacteria | Helicobacter cetorum | 138563 | 0.001709 | 7401 | 6375 | 3910 | 60.4 | 0.001243 |
| Bacteria | Helicobacter pylori | 210 | 4.451 | 19268562 | 15702849 | 2355117 | 385 | 0.06518 |
| Bacteria | Herbaspirillum rubrisubalbicans | 80842 | 0.0008103 | 3508 | 2754 | 2997 | 7.67 | 0.000332 |
| Bacteria | Herbaspirillum seropedicae | 964 | 0.0007169 | 3104 | 2489 | 3019 | 8.82 | 0.0003796 |
| Bacteria | Hydrogenophaga sp. PBL-H3 | 434010 | 0.0001862 | 806 | 806 | 3763 | 1.88 | 0.0009329 |
| Bacteria | Janthinobacterium lividum | 29581 | 0.00988 | 42775 | 22433 | 237250 | 2.59 | 0.04069 |
| Bacteria | Klebsiella aerogenes | 548 | 0.00106 | 4588 | 3718 | 1679 | 22.9 | 0.0001012 |

| | | | | | | | | |
|---|---|---|---|---|---|---|---|---|
| Bacteria | Klebsiella oxytoca | 571 | 0.0005301 | 2295 | 690 | 1988 | 17.4 | 0.0001511 |
| Bacteria | Klebsiella pneumoniae | 573 | 0.001243 | 5383 | 2059 | 2272 | 13.5 | 7.62E-05 |
| Bacteria | Kosakonia cowanii | 208223 | 0.0005218 | 2259 | 312 | 1083 | 22.3 | 0.0001787 |
| Bacteria | Leclercia adecarboxylata | 83655 | 0.0007327 | 3172 | 325 | 1074 | 31.6 | 0.0001055 |
| Bacteria | Lelliottia amnigena | 61646 | 0.0003102 | 1343 | 363 | 3210 | 3.71 | 0.0004666 |
| Bacteria | Marinobacter hydrocarbonoclasticus | 2743 | 0.00117 | 5066 | 2990 | 1034 | 59.2 | 0.0001313 |
| Bacteria | Moraxella osloensis | 34062 | 0.0007671 | 3321 | 1699 | 88753 | 1.36 | 0.009188 |
| Bacteria | Neisseria meningitidis | 487 | 9.08E-05 | 393 | 312 | 2988 | 1.84 | 0.0007454 |
| Bacteria | Neisseria mucosa | 488 | 0.0002455 | 1063 | 243 | 14293 | 1.21 | 0.004553 |
| Bacteria | Oblitimonas alkaliphila | 1697053 | 0.0006899 | 2987 | 1934 | 6107 | 5.5 | 0.001391 |
| Bacteria | Ochrobactrum anthropi | 529 | 0.0003458 | 1497 | 1428 | 6747 | 3.27 | 0.0008589 |
| Bacteria | Pandoraea pnomenusa | 93220 | 0.0007225 | 3128 | 2653 | 1469 | 13.4 | 0.0001756 |
| Bacteria | Pandoraea thiooxydans | 445709 | 0.0006174 | 2673 | 2673 | 1001 | 18.9 | 0.0002387 |
| Bacteria | Pantoea agglomerans | 549 | 0.000264 | 1143 | 753 | 7153 | 2.88 | 0.0007185 |
| Bacteria | Pantoea ananatis | 553 | 0.000173 | 749 | 433 | 1415 | 6.77 | 0.0001228 |
| Bacteria | Pantoea vagans | 470934 | 0.0007421 | 3213 | 649 | 2402 | 11.7 | 0.000177 |
| Bacteria | Paraburkholderia aromaticivorans | 2026199 | 0.0008128 | 3519 | 338 | 1377 | 13.8 | 9.49E-05 |
| Bacteria | Pectobacterium parmentieri | 1905730 | 0.0002522 | 1092 | 525 | 5521 | 3.59 | 0.0009178 |
| Bacteria | Proteus mirabilis | 584 | 0.0003056 | 1323 | 543 | 1232 | 4.92 | 0.0001031 |
| Bacteria | Pseudomonas aeruginosa | 287 | 0.02064 | 89352 | 49490 | 48320 | 12.1 | 0.001252 |
| Bacteria | Pseudomonas agarici | 46677 | 0.04295 | 185954 | 836 | 19401 | 63.5 | 0.004089 |
| Bacteria | Pseudomonas alcaligenes | 43263 | 0.00679 | 29399 | 914 | 10897 | 16.6 | 0.002987 |
| Bacteria | Pseudomonas alkylphenolica | 237609 | 0.03713 | 160747 | 68321 | 19723 | 55.2 | 0.002717 |
| Bacteria | Pseudomonas antarctica | 219572 | 0.1921 | 831876 | 11522 | 1637198 | 6.36 | 0.2201 |
| Bacteria | Pseudomonas arsenicoxydans | 702115 | 0.08846 | 382989 | 65170 | 62564 | 55.7 | 0.008105 |
| Bacteria | Pseudomonas azotoformans | 47878 | 0.2717 | 1176108 | 18963 | 120885 | 86.8 | 0.009002 |
| Bacteria | Pseudomonas balearica | 74829 | 0.005673 | 24560 | 14602 | 3571 | 40.1 | 0.000795 |
| Bacteria | Pseudomonas brassicacearum | 930166 | 0.06089 | 263601 | 61231 | 32826 | 58.6 | 0.002708 |
| Bacteria | Pseudomonas brenneri | 129817 | 0.05116 | 221510 | 451 | 31100 | 59.1 | 0.008357 |
| Bacteria | Pseudomonas cerasi | 1583341 | 0.002934 | 12704 | 12340 | 3930 | 28.5 | 0.003842 |
| Bacteria | Pseudomonas chlororaphis | 587753 | 0.1998 | 864887 | 371614 | 114017 | 53.3 | 0.002476 |
| Bacteria | Pseudomonas cichorii | 36746 | 0.09097 | 393840 | 2387 | 21013 | 136 | 0.003923 |
| Bacteria | Pseudomonas citronellolis | 53408 | 0.006228 | 26964 | 13369 | 4212 | 42.1 | 0.0005455 |

| | | | | | | | | |
|---|---|---|---|---|---|---|---|---|
| Bacteria | Pseudomonas coronafaciens | 53409 | 0.04762 | 206176 | 173401 | 16175 | 84.9 | 0.002668 |
| Bacteria | Pseudomonas corrugata | 47879 | 0.02694 | 116638 | 97954 | 14548 | 53.2 | 0.003099 |
| Bacteria | Pseudomonas entomophila | 312306 | 0.02352 | 101828 | 34500 | 63253 | 11 | 0.006227 |
| Bacteria | Pseudomonas extremaustralis | 359110 | 0.3731 | 1615381 | 4035 | 135401 | 123 | 0.03183 |
| Bacteria | Pseudomonas extremorientalis | 169669 | 0.06108 | 264458 | 612 | 29405 | 69.4 | 0.009077 |
| Bacteria | Pseudomonas fluorescens | 294 | 0.878 | 3801481 | 724305 | 564447 | 54.1 | 0.007131 |
| Bacteria | Pseudomonas fragi | 296 | 0.02145 | 92856 | 39212 | 45149 | 15.5 | 0.01098 |
| Bacteria | Pseudomonas frederiksbergensis | 104087 | 0.1379 | 597101 | 3210 | 68265 | 97 | 0.006536 |
| Bacteria | Pseudomonas fulva | 47880 | 0.009466 | 40982 | 1116 | 6523 | 33.6 | 0.001263 |
| Bacteria | Pseudomonas knackmussii | 65741 | 0.007414 | 32098 | 17568 | 5821 | 38.3 | 0.0008914 |
| Bacteria | Pseudomonas koreensis | 198620 | 0.09008 | 390000 | 82742 | 790992 | 4.86 | 0.06732 |
| Bacteria | Pseudomonas libanensis | 75588 | 0.04936 | 213706 | 442 | 41899 | 38.4 | 0.01044 |
| Bacteria | Pseudomonas lurida | 244566 | 0.01851 | 80142 | 10817 | 23457 | 32.6 | 0.0136 |
| Bacteria | Pseudomonas mandelii | 75612 | 0.02055 | 88959 | 12234 | 14135 | 56.4 | 0.004896 |
| Bacteria | Pseudomonas mediterranea | 183795 | 0.02252 | 97483 | 78076 | 12377 | 52.9 | 0.002499 |
| Bacteria | Pseudomonas mendocina | 300 | 0.02688 | 116356 | 50167 | 16252 | 47.8 | 0.0009549 |
| Bacteria | Pseudomonas migulae | 78543 | 0.0291 | 125989 | 650 | 20333 | 42.9 | 0.005539 |
| Bacteria | Pseudomonas monteilii | 76759 | 0.0179 | 77506 | 26508 | 15197 | 32.6 | 0.001868 |
| Bacteria | Pseudomonas mosselii | 78327 | 0.002737 | 11850 | 10733 | 1693 | 37.7 | 0.001363 |
| Bacteria | Pseudomonas mucidolens | 46679 | 0.06751 | 292292 | 292205 | 34352 | 65.6 | 0.007593 |
| Bacteria | Pseudomonas oleovorans | 301 | 0.004186 | 18122 | 9793 | 9089 | 15.8 | 0.002952 |
| Bacteria | Pseudomonas orientalis | 76758 | 0.1801 | 779761 | 323833 | 104544 | 57.7 | 0.00778 |
| Bacteria | Pseudomonas oryzihabitans | 47885 | 0.007657 | 33150 | 4098 | 6239 | 36.8 | 0.0005236 |
| Bacteria | Pseudomonas otitidis | 319939 | 0.007636 | 33058 | 17133 | 4338 | 47.4 | 0.0006351 |
| Bacteria | Pseudomonas parafulva | 157782 | 0.01674 | 72483 | 53982 | 8874 | 54 | 0.00165 |
| Bacteria | Pseudomonas plecoglossicida | 70775 | 0.009991 | 43257 | 40444 | 16557 | 17 | 0.003863 |
| Bacteria | Pseudomonas poae | 200451 | 0.1655 | 716698 | 169064 | 115574 | 53.3 | 0.0121 |
| Bacteria | Pseudomonas protegens | 380021 | 0.06976 | 302044 | 155202 | 32715 | 61.3 | 0.002518 |
| Bacteria | Pseudomonas psychrophila | 122355 | 0.04323 | 187176 | 186955 | 151724 | 9.83 | 0.03481 |
| Bacteria | Pseudomonas psychrotolerans | 237610 | 0.005161 | 22346 | 2258 | 3257 | 39.2 | 0.0004079 |
| Bacteria | Pseudomonas putida | 303 | 0.2186 | 946233 | 264149 | 1033238 | 8.83 | 0.01301 |
| Bacteria | Pseudomonas reinekei | 395598 | 0.03591 | 155461 | 259 | 20693 | 55.3 | 0.004592 |

| | | | | | | | | |
|---|---|---|---|---|---|---|---|---|
| Bacteria | Pseudomonas rhodesiae | 76760 | 0.1169 | 505941 | 304834 | 58767 | 83.2 | 0.01079 |
| Bacteria | Pseudomonas simiae | 321846 | 0.1017 | 440503 | 365445 | 50288 | 82.2 | 0.01073 |
| Bacteria | Pseudomonas sp. | 306 | 0.01243 | 53814 | 39447 | 6735 | 107 | 0.00892 |
| Bacteria | Pseudomonas sp. Lz4W | 1206777 | 0.003745 | 16214 | 632 | 10061 | 13.8 | 0.01384 |
| Bacteria | Pseudomonas sp. R32 | 1573704 | 0.0248 | 107362 | 470 | 10467 | 59.9 | 0.002555 |
| Bacteria | Pseudomonas synxantha | 47883 | 0.1725 | 746694 | 400981 | 110632 | 55.1 | 0.008012 |
| Bacteria | Pseudomonas taetrolens | 47884 | 0.03183 | 137812 | 137036 | 21551 | 45.2 | 0.005104 |
| Bacteria | Pseudomonas thivervalensis | 86265 | 0.02051 | 88795 | 77794 | 11878 | 59.2 | 0.002475 |
| Bacteria | Pseudomonas tolaasii | 29442 | 0.08528 | 369239 | 2752 | 38442 | 81.1 | 0.00836 |
| Bacteria | Pseudomonas trivialis | 200450 | 0.1507 | 652298 | 1281 | 73187 | 71.2 | 0.01013 |
| Bacteria | Pseudomonas umsongensis | 198618 | 0.03927 | 170015 | 127075 | 26295 | 48.5 | 0.003667 |
| Bacteria | Pseudomonas veronii | 76761 | 1.041 | 4506550 | 1675097 | 322936 | 166 | 0.06806 |
| Bacteria | Pseudomonas viridiflava | 33069 | 0.05019 | 217298 | 1752 | 16078 | 88.9 | 0.003072 |
| Bacteria | Pseudomonas yamanorum | 515393 | 0.1266 | 548151 | 159430 | 519052 | 10.1 | 0.05296 |
| Bacteria | Psychrobacter cryohalolentis | 330922 | 0.0002282 | 988 | 987 | 9564 | 1.45 | 0.00677 |
| Bacteria | Rahnella aquatilis | 34038 | 0.001588 | 6876 | 1136 | 25982 | 2.51 | 0.003987 |
| Bacteria | Ralstonia insidiosa | 190721 | 0.0008789 | 3805 | 1027 | 6953 | 4.43 | 0.0008925 |
| Bacteria | Ralstonia mannitolilytica | 105219 | 0.0001377 | 596 | 441 | 5428 | 2.18 | 0.00144 |
| Bacteria | Ralstonia pickettii | 329 | 0.0006393 | 2768 | 585 | 68577 | 1.48 | 0.004922 |
| Bacteria | Ralstonia solanacearum | 305 | 0.0007544 | 3266 | 1588 | 2701 | 8.93 | 9.58E-05 |
| Bacteria | Raoultella terrigena | 577 | 0.0004414 | 1911 | 260 | 2312 | 6.81 | 0.0004435 |
| Bacteria | Rhizobacter gummiphilus | 946333 | 0.0003042 | 1317 | 1317 | 1797 | 5.07 | 0.0003061 |
| Bacteria | Rhizobium leguminosarum | 384 | 0.001125 | 4869 | 1968 | 1680 | 21.3 | 2.94E-05 |
| Bacteria | Rhodopseudomonas palustris | 1076 | 0.0008093 | 3504 | 957 | 8805 | 2.87 | 0.0002211 |
| Bacteria | Salmonella enterica | 28901 | 0.001596 | 6909 | 3166 | 4315 | 16.8 | 6.37E-05 |
| Bacteria | Schaalia odontolytica | 1660 | 0.0004472 | 1936 | 379 | 29135 | 1.21 | 0.007272 |
| Bacteria | Serratia fonticola | 47917 | 0.0007916 | 3427 | 792 | 2440 | 18.9 | 0.0001701 |
| Bacteria | Serratia liquefaciens | 614 | 0.0004435 | 1920 | 747 | 1057 | 11.7 | 0.0001954 |
| Bacteria | Serratia marcescens | 615 | 0.001846 | 7994 | 4655 | 7019 | 12.5 | 0.0002348 |
| Bacteria | Serratia plymuthica | 82996 | 0.0009239 | 4000 | 2340 | 4044 | 8.23 | 0.0002949 |
| Bacteria | Serratia symbiotica | 138074 | 0.0003499 | 1515 | 347 | 1125 | 17.5 | 0.0001327 |
| Bacteria | Sphingobium yanoikuyae | 13690 | 0.0003698 | 1601 | 310 | 23986 | 1.3 | 0.002369 |
| Bacteria | Sphingopyxis macrogoltabida | 33050 | 0.0003137 | 1358 | 329 | 2654 | 1.83 | 0.0003327 |
| Bacteria | Stenotrophomonas acidaminiphila | 128780 | 0.0001989 | 861 | 397 | 10072 | 1.49 | 0.001281 |
| Bacteria | Stenotrophomonas maltophilia | 40324 | 0.001432 | 6198 | 2942 | 72311 | 1.84 | 0.001447 |

| | | | | | | | | |
|---|---|---|---|---|---|---|---|---|
| Bacteria | Thalassolituus oleivorans | 187493 | 0.0002448 | 1060 | 749 | 10410 | 1.62 | 0.00183 |
| Bacteria | uncultured Pseudomonas sp. | 114707 | 0.001151 | 4983 | 677 | 2675 | 10.4 | 0.00773 |
| Bacteria | Variovorax paradoxus | 34073 | 0.002156 | 9333 | 917 | 27495 | 2.95 | 0.0008695 |
| Bacteria | Vibrio cholerae | 666 | 0.0004414 | 1911 | 1106 | 1275 | 15.6 | 7.16E-05 |
| Bacteria | Xanthomonas campestris | 339 | 0.0004855 | 2102 | 325 | 15764 | 2.84 | 0.001191 |
| Bacteria | Xanthomonas citri | 346 | 0.0007327 | 3172 | 713 | 1737 | 13.3 | 0.0001183 |
| Bacteria | Yersinia enterocolitica | 630 | 0.0001213 | 525 | 308 | 1592 | 2.63 | 0.0001446 |
| Bacteria | Yersinia ruckeri | 29486 | 0.0001086 | 470 | 335 | 1572 | 4.79 | 0.0002772 |
| Bacteria | uncultured prokaryote | 198431 | 0.002813 | 12180 | 1456 | 5410 | 21.5 | 0.0003763 |
| Eukaryota | uncultured eukaryote | 100272 | 0.0007911 | 3425 | 735 | 5634 | 4.75 | 0.0006768 |
| Fungi | Antarctomyces pellizariae | 1955577 | 0.001575 | 6821 | 6821 | 1056 | 190 | 0.04621 |
| Fungi | Apiotrichum porosum | 105984 | 0.00244 | 10563 | 373 | 1111 | 54.2 | 8.57E-05 |
| Fungi | Aspergillus nidulans | 162425 | 0.00148 | 6407 | 580 | 135510 | 1.16 | 0.004575 |
| Fungi | Aspergillus niger | 5061 | 9.03E-05 | 391 | 222 | 1864 | 2.57 | 8.33E-05 |
| Fungi | Botrytis cinerea | 40559 | 0.0004633 | 2006 | 1557 | 1524 | 8.21 | 3.42E-05 |
| Fungi | Leucosporidium scottii | 5278 | 0.006938 | 30038 | 5754 | 1718 | 185 | 0.001616 |
| Fungi | Leucosporidium sp. AY30 | 662878 | 0.04991 | 216072 | 654 | 2463 | 2070 | 0.894 |
| Fungi | piotrichum mycotoxinovorans | 252803 | 0.003578 | 15492 | 9198 | 1413 | 69.8 | 4.32E-05 |
| Fungi | Pseudogymnoascus destructans | 655981 | 0.2999 | 1298527 | 60481 | 473667 | 37.5 | 0.03685 |
| Fungi | Pseudogymnoascus pannorum | 79858 | 0.1009 | 437040 | 205 | 4830 | 1830 | 0.1145 |
| Fungi | Pseudogymnoascus verrucosus | 342668 | 0.1819 | 787723 | 569 | 405670 | 21.7 | 0.03034 |
| Fungi | Saccharomyces cerevisiae | 4932 | 0.000212 | 918 | 641 | 15459 | 1.24 | 0.0006803 |
| Fungi | Talaromyces rugulosus | 121627 | 0.0003458 | 1497 | 1134 | 4532 | 3.34 | 0.0001338 |
| Fungi | Malassezia globosa | 76773 | 0.0001894 | 820 | 746 | 6336 | 1.88 | 0.0007186 |
| Fungi | Malassezia restricta | 76775 | 0.002037 | 8820 | 8477 | 54633 | 2.27 | 0.006595 |
| Fungi | Phaffia rhodozyma | 264483 | 0.001012 | 4382 | 652 | 1887 | 16.3 | 0.0001045 |
| Fungi | Phenoliferia glacialis | 418497 | 0.003383 | 14645 | 13461 | 1056 | 72.5 | 0.2884 |
| Fungi | Rhodotorula toruloides | 5286 | 0.04413 | 191057 | 127887 | 14868 | 77.8 | 0.0007132 |
| Fungi | Schizophyllum commune | 5334 | 0.002495 | 10804 | 249 | 1185 | 53.5 | 6.57E-05 |
| Fungi | Sporisorium reilianum | 72558 | 0.002726 | 11801 | 8808 | 1294 | 44.4 | 5.89E-05 |
| Fungi | Anthracocystis flocculosa | 84751 | 0.00434 | 18791 | 253 | 2257 | 54.6 | 0.0001896 |
| Fungi | uncultured fungus | 175245 | 0.001476 | 6392 | 1470 | 13970 | 3.57 | 0.0008285 |
| Non-applicable | BMMF2 DNA sequence | 2502152 | 0.0001779 | 770 | 641 | 3644 | 6.05 | 0.05802 |
| Non-applicable | synthetic construct | 32630 | 0.00378 | 16365 | 9071 | 28345 | 47 | 0.001601 |
| Non-applicable | uncultured organism | 155900 | 0.0005197 | 2250 | 412 | 7743 | 7.9 | 0.0003169 |
| Plantae | Aegilops tauschii | 37682 | 0.01224 | 53003 | 1372 | 324531 | 1.79 | 0.006682 |

| | | | | | | | | |
|---|---|---|---|---|---|---|---|---|
| Plantae | Digitaria exilis | 1010633 | 0.000306 | 1325 | 1325 | 3340 | 2.14 | 7.82E-06 |
| Plantae | Hordeum vulgare | 4513 | 0.009172 | 39709 | 4512 | 227095 | 2.06 | 0.004067 |
| Plantae | Oryza sativa | 4530 | 0.0004132 | 1789 | 627 | 4047 | 2.86 | 1.13E-05 |
| Plantae | Secale cereale | 4550 | 0.001784 | 7723 | 1086 | 19502 | 4.01 | 0.02282 |
| Plantae | Triticum aestivum | 4565 | 0.337 | 1458842 | 445502 | 3735086 | 4.77 | 0.07944 |
| Plantae | Triticum monococcum | 4568 | 0.03776 | 163467 | 17938 | 305992 | 6.27 | 0.1575 |
| Plantae | Triticum turgidum | 4571 | 0.02711 | 117387 | 2202 | 297714 | 3.84 | 0.1045 |
| Plantae | Triticum urartu | 4572 | 0.01042 | 45106 | 1315 | 91006 | 4.79 | 0.1283 |
| Plantae | Zea mays | 4577 | 0.000231 | 1000 | 203 | 1002 | 3.54 | 1.23E-05 |
| Virus | uncultured Caudovirales phage | 2100421 | 0.003801 | 16458 | 10444 | 5373 | 43 | 0.00011 |
